# Early and current environments exert distinct effects on immune function in the Orang Asli

**DOI:** 10.64898/2026.07.24.740585

**Authors:** Layla Brassington, Audrey M. Arner, Grace Rodenberg, Nicholas Ryan, Diane Song, Tan Bee Ting A/P Tan Boon Huat, Virginia B. Kraus, Janet L. Huebner, Izandis bin Mohd Sayed, Yvonne Ai Lian Lim, Vivek V. Venkataraman, Ian J. Wallace, Thomas S. Kraft, Amanda J. Lea

## Abstract

Humans evolved in environments characterized by subsistence foraging and hunting, high levels of physical activity, and regular and diverse pathogen exposure. Industrialization has rapidly altered these environments, leading to increased metabolic and inflammatory disease risk. These observations have motivated studies of how industrialization shapes immune function, but whether industrialization alters immune biology through developmental embedding of early-life environments or ongoing plastic responses to current conditions remains poorly understood. To address this gap, we worked with the Orang Asli—the Indigenous peoples of Peninsular Malaysia—who currently span a gradient from subsistence horticulture, foraging, and hunting to industrialized, urban environments with substantial inter-individual variation in life course experiences. Leveraging continuous measures of both early-life and current lifestyle, we found that current lifestyle exerts stronger effects on adult immune gene expression than early-life conditions, impacting 1,428 as compared to 223 genes, respectively. Early-life associated genes were enriched for pathways regulating adaptive immunity, particularly T cell development and differentiation, consistent with higher predicted T cell abundance and circulating IL-8 in urban-born individuals. In contrast, current exposure to urban, industrialized conditions was linked to innate immune and inflammatory activation, including dendritic cell abundance, metabolic pathway upregulation, and elevated CRP. Finally, consistent with lower rates of immune disorders in non-industrial settings, current exposure to this lifestyle was associated with up-regulation of genes involved in Th1/Th2 differentiation. Together, these results emphasize that industrialized immune profiles reflect both early-life developmental embedding and ongoing environmental responses, highlighting the importance of considering exposures across the life course.

**Significance statement:** Humans evolved in environments with high levels of physical activity, subsistence-based diets, and constant pathogen exposure. Rapid industrialization has dramatically changed these conditions, coinciding with rising rates of non-communicable diseases with inflammatory underpinnings. Our study with the Orang Asli, the Indigenous peoples of Peninsular Malaysia, tests how exposure to industrialization across the life course shapes immune variation. We find that early-life environments leave lasting marks on adaptive immune function particularly T cells, while industrialized adult lifestyle influences innate immune activity and inflammation. These results clarify how different stages of life contribute to immune remodeling and point to a mechanistic link between industrialization and disease risk, highlighting the importance of both early-life and current environments in shaping later life health.

## Introduction

For most of human evolutionary history, immune systems were shaped by constant exposure to diverse pathogens in resource-limited conditions, forcing trade-offs between investment in immune function and other necessary biological processes [1–6]. Over the last few generations, however, many populations have transitioned rapidly from subsistence-based, non-industrial livelihoods to urban, market-integrated, and industrialized environments, reshaping diet, activity patterns, and pathogen exposures in ways that fundamentally alter immune investment and biology [7,8]. These transitions have also coincided with sharp rises in noncommunicable diseases such as obesity, type 2 diabetes, and cardiovascular disease [9–16], all conditions with a strong inflammatory component [8,17,18]; together, these observations have sparked interest in how urban, industrialized lifestyles impact immune function [7,8,19–21]. For example, comparisons between adults living in urban versus rural locations within both Senegal and Indonesia have revealed differences in T helper cell polarization, memory cell pools, and overall shifts toward innate immune activation in urban residents [22–24]. Similarly, experimental, short-term diet interventions from traditional to Western foods in a Tanzanian cohort were associated with coordinated changes in the plasma proteome, metabolome, and whole-blood transcriptome, generally pointing to up-regulation of innate inflammatory processes in individuals consuming a Western diet [25].

While impactful, much of this previous work has relied on binary comparisons of “urban” and “rural” individuals. These designs limit inference in two important ways: they reduce lifestyle to discrete categories despite substantial within-population variation, and because early-life and current environments are typically aligned, they cannot determine whether immune differences reflect developmental embedding or ongoing adult plasticity. This creates an important gap because early-life environments—including nutritional conditions, psychosocial stress, and microbial exposures during the pre- and peri-natal period—are well known to influence thymic output, B and T cell repertoires, and gene expression programs within immune cells [26–29]. For example, experimental early-life microbial exposure in mice alters the CD8 T cell compartment, promoting the preferential expansion of highly responsive fetal-derived CD8 T cells that persist into adulthood and enhance later life protection against intracellular pathogens [30]. Similarly, in a longitudinal human cohort study in the Philippines, greater exposure to animals and infection in infancy predicted lower inflammation (as measured by CRP) in adulthood, emphasizing the long reach of early-life experiences [27,31]. Together, these studies and related findings [32–36] demonstrate that early-life exposures, especially those that occur during the “window of opportunity” for immune priming [26], can durably structure immune cell composition, regulatory architecture, and inflammatory biology in ways that persist into adulthood.

Nevertheless, despite an increasing appreciation that early-life environments matter, it remains unresolved whether immune differences between “urban” and “rural” adults primarily reflect developmental embedding or ongoing responses to current environments. This is partly because distinguishing the effects of early-life versus later-life environments requires working with populations where these variables are separable. To do so, we drew on our ongoing relationships with a population that experiences substantial within and between individual variation in industrialized lifestyle exposures—the Indigenous peoples of Peninsular Malaysia, collectively known as the Orang Asli [37,38]. The Orang Asli make up <1% of the Peninsular population, and include 19 ethnolinguistic groups divided into three major subgroups (Proto-Malay, Senoi, and Negrito). Over the last half-century, all groups have experienced varying degrees of lifestyle change, which our research team and others have linked to reduced physical activity, sleep alterations, increased access to market foods, and rising cardiometabolic disease risk in more urban, industrialized communities [9,39–41]. These lifestyle shifts are driven in part by deforestation of traditional lands and expansion of plantation agriculture [42–44], as well as efforts by the Malaysian government to assimilate the Orang Asli into mainstream culture, including through new economic and resettlement schemes focused on “modern” facilities [45–50]. Due to the heterogeneity of these forces, some communities still live in remote, rainforest camps and rely heavily on natural resources [38,51], while others are fully integrated into the cash economy; importantly, because the pace of market-integration and acculturation has varied widely, there is substantial inter-individual variation in the timing of urban lifestyle exposures [50,52,53]. Finally, a unique social movement among some Orang Asli groups, called “back to roots”, is prompting individuals that are dissatisfied with urban living (e.g., because of overcrowding, limited economic opportunities, and government pressures to assimilate) to return to their traditional lands. Taken together, the Orang Asli thus represent a unique participant population, with individuals having experienced a high degree of within-lifetime lifestyle variation—including both non-industrial to industrialized shifts that are playing out globally, as well as reverse transitions that are extremely rare.

Leveraging this natural variation, we worked with Orang Asli adults to collect: (1) questionnaire data about both early-life and current lifestyle exposures; (2) quantitative measures of individual factors hypothesized to mediate the relationship between lifestyle and immune function, specifically adiposity [54], physical activity measured via accelerometry [39], eosinophil and IgE levels reflecting helminth exposure [55], and indices of both traditional and market diet [8]; and (3) comprehensive immune outcomes, including immune cell composition (n=922), transcriptomics (n=922), and cytokines and inflammatory biomarkers (n=317). Integrating these datasets, we tested three main hypotheses: (1) early-life conditions have lasting effects on adaptive immune composition and gene expression programs (especially for T cells); (2) current urban lifestyles primarily impact innate immune cells and alter inflammatory signaling; and (3) up-regulation of proinflammatory processes in urban residents is proximately driven by low levels of physical activity, consumption of ultra-processed foods, and central adiposity. By jointly analyzing continuous early-life and current lifestyle measures with physiological, cellular, and molecular immune phenotypes, this study aims to disentangle developmental embedding from adult plasticity and to clarify the pathways through which industrialized environments shape immune and metabolic health.

## Results

### Variation in early-life and current environments among the Orang Asli

To disentangle early-life and current lifestyle effects on later life immune traits, we drew on questionnaire and biological datasets for 922 adult participants (spanning ages 18 to 91 years, and including 577 females and 345 males, respectively) (SI Figure 1). These data were collected as part of the Orang Asli Health and Lifeways Project (OA HeLP), an ongoing study of Orang Asli culture, health, and well-being in the face of lifestyle change [37] (Figure 1A-B). To operationalize our life course approach, we distilled questionnaires (Table S1) about early-life and current environments into a composite “early-life lifestyle score” and “current lifestyle score” using approaches derived from the literature [56,57]; we have since validated these scores and their effects on adult cardiometabolic health in Orang Asli [15,41]. Briefly, the early-life lifestyle score includes information about parental occupation (e.g., engaging in foraging, horticulture, or fishing versus wage labor), housing (e.g., living in a traditional bamboo house versus one made of wood or brick), access to market-derived foods and possessions (e.g., salt, sugar, oil, phones, televisions), and access to urban infrastructure (e.g., piped water supply, electricity, roads, sewage) during childhood. The current lifestyle score averages most of the same components across individuals living in a given location, and can be thought of as capturing a parallel axis of variation for the contemporary environment [9,41]. For both scores, higher values are associated with more urban, industrialized lifestyles. Additional information about score construction is provided in Supplementary Materials.

**Figure 1:**
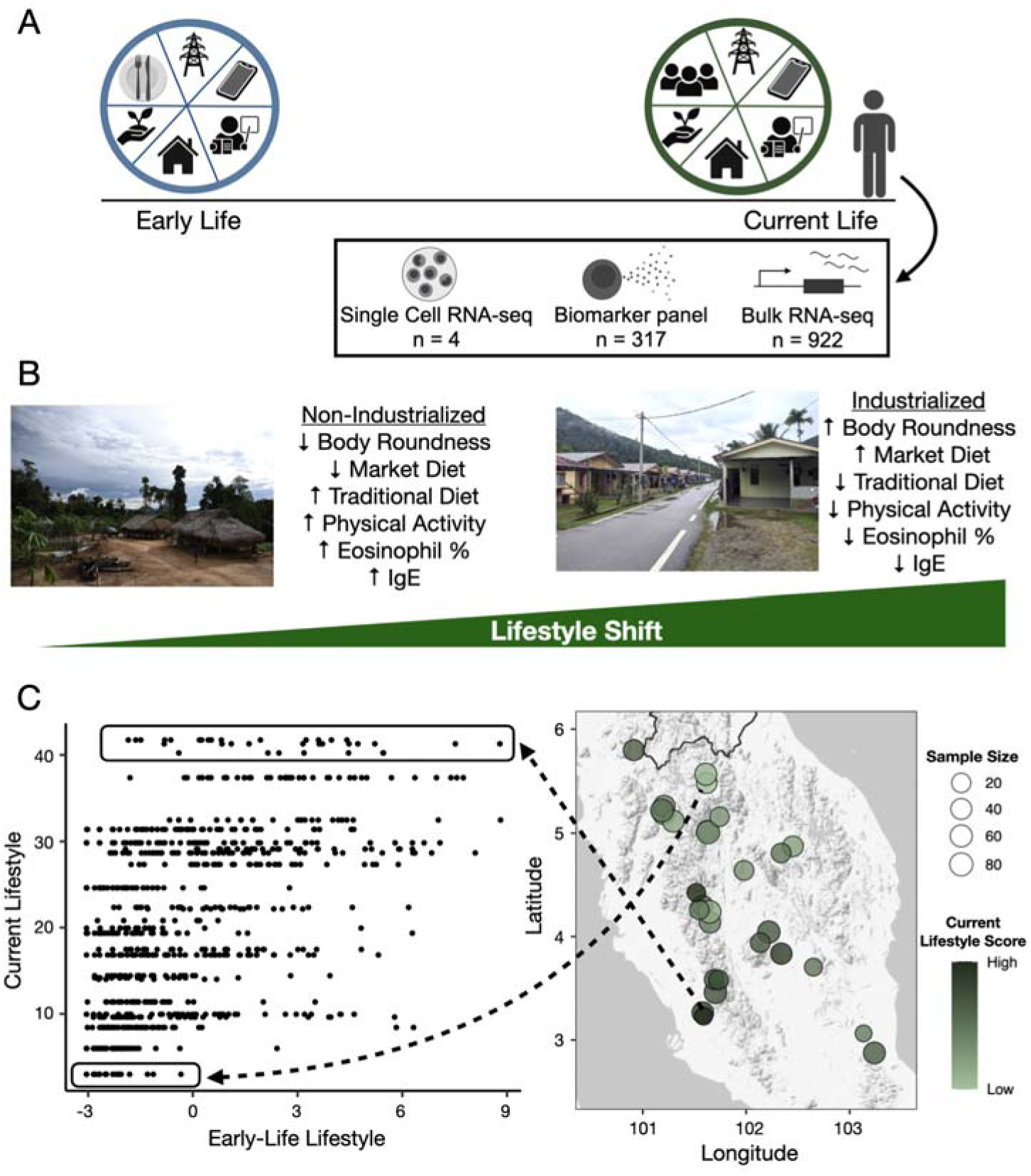
Study design and overview of early-life and current lifestyle variation. (A) Study design. Adult participants were interviewed about their early-life and current lifestyle experiences, at which time biological samples were also collected. Survey responses were distilled into a continuous, composite “early-life lifestyle score” and “current lifestyle score” using previously described approaches [15,41]. The domains of information included in each composite score are noted in the boxes. (B) Individual, measured environmental components that have been linked to the industrialization gradient in previous work [16,39] and that correlate with the current lifestyle score in this study (all p<0.05; see “Composite lifestyle score captures broader transcriptomic effects than individual lifestyle components” for further details). Photos show opposite ends of the lifestyle spectrum from non-industrial, subsistence-based communities to urban, industrialized locations. (C) Left panel: relationship between early-life lifestyle score (x-axis) and current lifestyle score (y-axis). Right panel: participant communities. Point size corresponds to the number of participants, and color reflects the mean current lifestyle score. Arrows highlight two example locations and participant’s within-lifetime lifestyle variation.

Consistent with previous work with Orang Asli [15], we observe considerable lifestyle variation both between and within individuals (Figure 1C), with a moderate correlation between early-life and current environments (Spearman correlation, R² = 0.26, p = 8.42x10^-63^). For example, some individuals (lower left of Figure 1C) have lived their entire lives in non-industrial, subsistence-based environments, while others who grew up in these environments now live in urban, industrialized communities (upper left of Figure 1C). Notably, this population also includes individuals who grew up in urban environments but have since returned to traditional lands (lower right of Figure 1C). Together, these bidirectional life course trajectories make the Orang Asli uniquely informative for disentangling the independent and combined effects of early-life and current environments.

### Early-life environments have targeted effects on adaptive immune cell types, while current environments broadly shape immune cell composition

Urbanization may influence immune function in part by reshaping the composition of circulating immune cells. To test this possibility, we quantified the abundance of major immune cell populations in peripheral blood mononuclear cells (PBMCs), a circulating compartment that integrates environmental exposures and contributes to cardiometabolic and inflammatory disease processes [58–63]. Using single-cell RNA-seq (scRNA-seq) data from 16,237 cells for four Orang Asli individuals, we defined marker sets for six major immune cell types and applied these to bulk PBMC RNA-seq profiles across the full cohort (n = 922) using standard pipelines [64,65] (Figure 2B, SI Figure 2). As expected, predicted immune cell proportions were correlated with both lymphocytes (r = 0.32, p = 1.30×10^-23^) and monocytes (r = 0.09, p = 0.006) measured via 5-part white blood cell differentials. The distribution of predicted cell types fell within expected physiological ranges for healthy individuals [66] (SI Figure 2).

**Figure 2:**
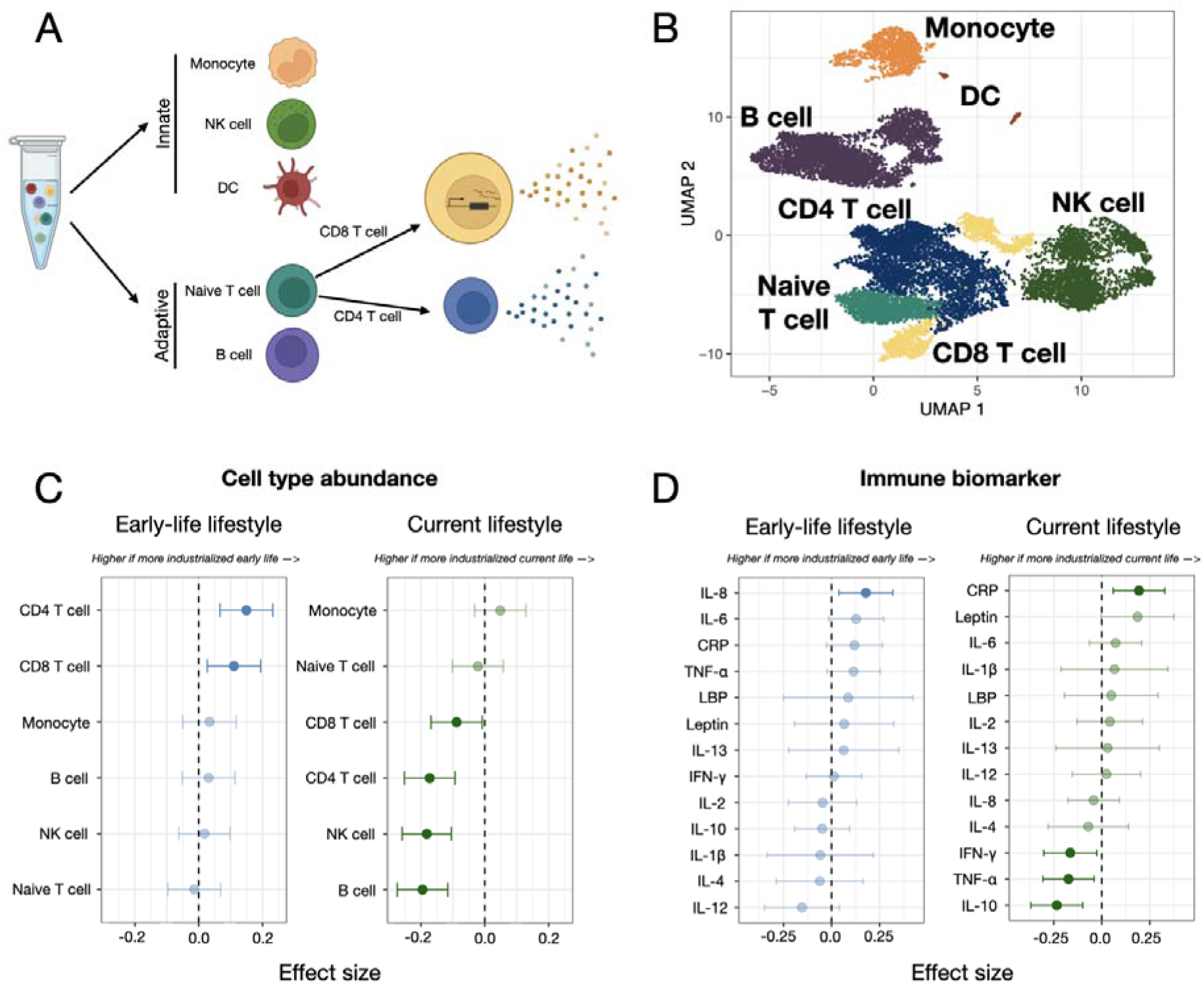
Lifestyle effects on immune cell composition and biomarkers. (A) Immune cell types included in deconvolution analyses. Peripheral blood mononuclear cells (PBMCs) represent a heterogeneous mixture of immune cell types spanning both innate and adaptive compartments. Each cell type exhibits a distinct transcriptomic profile that contributes to variation in cytokine abundance. Abbreviations: DC, dendritic cell; NK, natural killer. (B) Uniform manifold approximation and projection (UMAP) embedding of Orang Asli single-cell RNA-seq data used as the reference for bulk RNA-seq deconvolution (n = 4), with cells colored by their annotated immune cell type. (C-D) Associations between early-life and current lifestyle scores and immune traits. Points show model effect estimates from regressions of relative cell type proportions (C) or circulating biomarker abundance (D) on each score. Higher values reflect higher predicted abundance in individuals with more industrialized lifestyles. All predictor and response variables were standardized (to a mean of zero and variance of 1) such that effect sizes are comparable. Point opacity indicates statistical significance after false discovery rate correction, with more opaque points representing associations with adjusted p < 0.1. Abbreviations: IL, interleukin; TNF-α, tumor necrosis factor alpha; CRP, C-reactive protein; IFN-γ, interferon gamma; LBP, Lipopolysaccharide-binding protein.

Using the predicted PBMC cell type proportions, we tested whether early-life and current lifestyle were associated with adult immune cell composition, controlling for age and sex. Early-life lifestyle was significantly associated with both CD4 and CD8 T cell proportions, which were consistently higher in individuals who grew up in more urban, industrialized environments (CD4 β = 0.0021, adjusted p = 0.0009; CD8 β = 0.0019, adjusted p = 0.018). Because T cell compartments and adaptive immune repertoires are shaped during early antigen exposure and thymic development [67,68], these results are consistent with developmental calibration of lymphocyte composition that persists into adulthood. In contrast, current lifestyle was most strongly associated with natural killer (NK) cell abundance (β = -0.004, adjusted p = 1.15x10^-5^), which was higher in individuals who grew up in more non-industrial settings (Figure 2C). NK cells are cytotoxic innate lymphocytes that play a key role in early immune surveillance and inflammatory signaling, with activity and abundance known to respond to recent infections and other contemporary environmental factors [69,70].

### Non-industrial environments promote immune activation, while urban environments promote systemic inflammation

Differences in immune cell proportions reveal how environments shape immune architecture, but they do not necessarily reflect changes in cell function. To capture these effects, we measured 13 circulating cytokines and inflammatory biomarkers, which provide a readout of key signaling molecules produced by immune cells in a subset of individuals (n = 317; Figure 2A). Individuals exposed to more urban early life conditions exhibited higher levels of IL-8 (β = 0.18, adjusted p = 0.073; Figure 2D), a chemokine involved in leukocyte recruitment and CD8 T cell differentiation [71–73]. In contrast, current lifestyle showed broader associations with circulating inflammatory markers (Figure 2D). Individuals living more non-industrial lifestyles exhibited higher IFN-γ (primarily produced by NK cells) and TNF-α levels (IFN-γ β = -0.16, adjusted p = 0.10; TNF-α β = -0.17, adjusted p = 0.07, respectively). These cytokines are central orchestrators of cell-mediated immune responses and are often co-regulated during pathogen-associated immune activation [74,75]. Individuals living more non-industrial lifestyles exhibited higher levels of IL-10 (β = -0.23, adjusted p = 0.008), an immunoregulatory cytokine that modulates responses to prevent inflammatory and autoimmune pathologies. In contrast, CRP—a major biomarker of systemic inflammation produced in the liver—was elevated among individuals living more urban, industrialized lifestyles (β = 0.19, adjusted p = 0.037), consistent with the proinflammatory factors (e.g., ultra processed foods, inactivity, central adiposity) known to be common in these environments.

### Environmental variation across the life course shapes adult immune gene expression

Environmental exposures are known to shape immune function through changes in gene regulation [76–78]. Thus, we used our measures of early-life and current lifestyle to test how they independently shape adult PBMC gene expression (n = 9,993 expressed genes, 922 individuals; see Methods). Current lifestyle exerted substantially broader transcriptomic impacts than early-life conditions, associating with 1,428 compared to 223 genes, respectively (adjusted p < 0.1; Figure 3A). Across all genes, the effects of current lifestyle were generally larger in magnitude than the effects of early-life lifestyle (t-test comparing absolute effect sizes, p = 9x10^-4^; SI Figure 4). Nevertheless, the set of genes targeted by early-life and current environments were more overlapping than expected by chance (n = 65 shared genes; Fisher’s exact test, log (OR) = 1.34, p = 8.94x10^-9^; SI Figure 4). Overlapping genes were generally associated with core, rather than immune-related, cellular processes (e.g., “response to unfolded protein”; see Table S2). Among the overlapping set, 64 of 65 genes exhibited opposing effects, suggesting early-life and current environments tune shared molecular pathways in contrasting ways rather than reinforcing one another (*EMP1* was the only gene to not follow this pattern).

**Figure 3:**
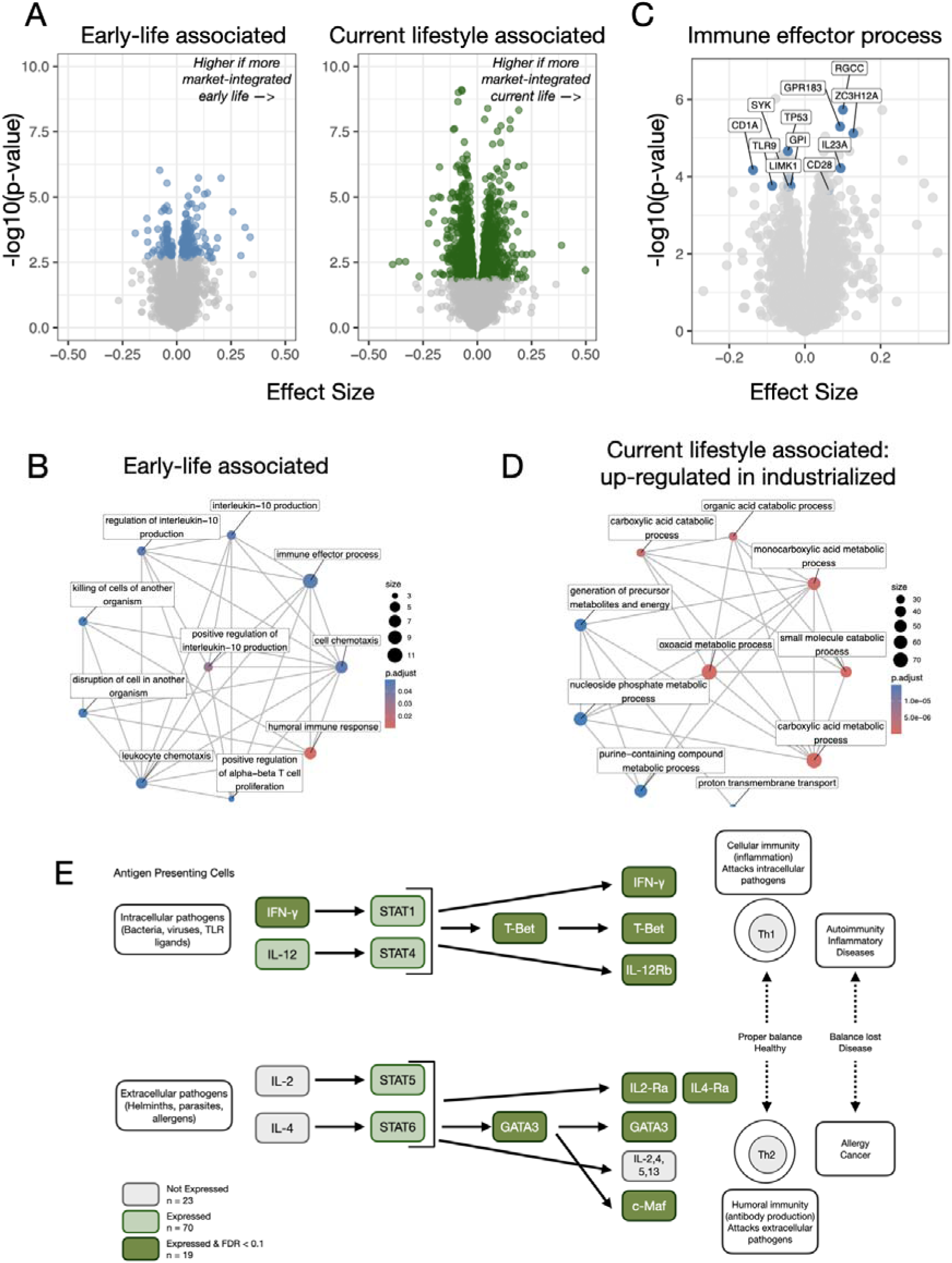
Effects of early-life and current lifestyle on gene expression in PBMCs. (A) Overlap of genes significantly associated with early-life and current lifestyle scores (adjusted p < 0.1). Volcano plots showing the effect size and statistical significance of associations between lifestyle scores and gene expression (GE). Specifically, the x-axis shows the effect size for each gene representing the effect of early-life or current lifestyle on gene expression, and the y-axis shows the −log10(p-value). (B&D) Gene ontology (GO) enrichment analysis of genes associated with environmental scores. Enriched GO terms are shown for genes associated with early-life lifestyle (B) and current lifestyle (D). Warmer colors indicate a lower FDR adjusted p-value and the size of the point represents the number of genes that overlap between the gene set of interest and the test gene set. (C) Volcano plot of genes belonging to the GO term “immune effector process,” which was significantly enriched among early-life-associated genes. Genes in the pathway are highlighted in blue, all other genes are grey. (E) Overlap between current non-industrial genes and the T helper cell differentiation pathway. Simplified version of the full KEGG pathway shown in SI Figure 8, displayed for improved readability. Colors indicate whether a gene was present in the KEGG pathway but not detected in our dataset (grey), present in both datasets but not significantly associated with current non-industrial environments (light green), or present in both datasets and significantly associated with current non-industrial environments (FDR adjusted p < 0.1; dark green).

**Figure 4:**
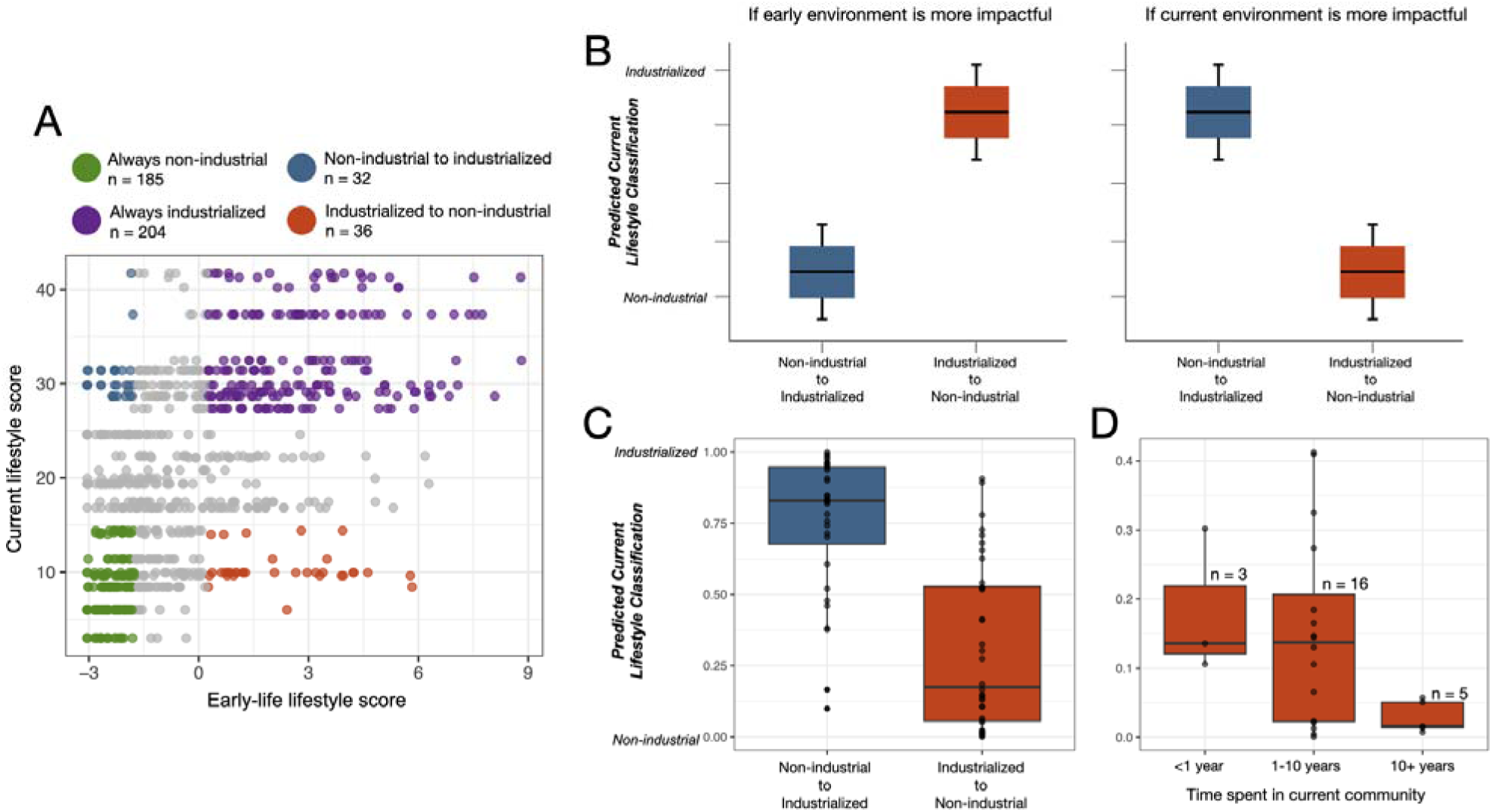
Within-lifetime transitions distinguish early-life embedding from plasticity in adult gene expression. (A) Distribution of individuals across early-life (x-axis) and current lifestyle (y-axis) scores. Points are colored according to lifestyle bin classification. Grey points indicate individuals who did not meet the threshold criteria for assignment to a lifestyle bin, which focused on environmental extremes (top and bottom ⅓ of scores for early-life and current lifestyle). (B) Conceptual overview of elastic net classifier predictions under alternative hypotheses in which adult gene expression reflects either early-life or current environmental conditions. (C) Confidence in elastic-net predicted bins for individuals who transitioned between environments during their lifetime. Confidence predictions range from 0 to 1, where values closer to 1 indicate gene expression profiles more similar to individuals who have lived in urban, industrialized environments for their entire life and values closer to 0 indicate profiles more similar to individuals that have lived in non-industrial, subsistence-based environments their entire life. (D) Relationship between predicted lifestyle assignment and estimated time spent in the current environment among individuals who transitioned from urban, industrialized to non-industrial, subsistence-based environments and received an elastic-net predicted score < 0.5. We note that we did not perform complementary analyses for individuals in the “non-industrial to industrialized" bin because there was no heterogeneity in the relevant question about timing (i.e., 100% of individuals underwent this lifestyle transition >10 years ago).

Genes associated with early-life lifestyle (in any direction, n = 223 genes) were enriched for adaptive immune processes, including “regulation of IL-10 production” and “immune effector process” (Table S2; Figure 3B). These genes included major immune orchestrators such as *CXCL8*, which encodes IL-8 and plays a key role in T cell differentiation as well as immune cell recruitment to sites of inflammation, infection, or injury [79–81] (Figure 2C–D). Genes driving enrichment of the top pathway (“immune effector process”) were up-regulated in both early-life non-industrial and urban environments, indicating that this enrichment reflects bidirectional transcriptional differences (Figure 3C). To identify upstream regulatory programs, we tested early-life associated genes for enrichment within the targets of individual transcription factors (TF) using the TRRUST v2 database [82], identifying 21 significantly enriched TFs (Fisher’s exact test, adjusted p < 0.1; SI Figure 6; Table S3). Among the top signals were *NF-*κ*B* (NFKB1/RELA complex; log2(OR) = 4.95, adjusted p = 3.54×10 ¹¹) and *SP1* (log2(OR) = 3.19, adjusted p = 8.52×10□¹□), both of which are major regulators of cellular responses to inflammatory and environmental stimuli [83,84].

Genes associated with current lifestyle showed distinct functional patterns depending on effect direction. Genes upregulated in non-industrial environments (n = 793 genes) were not significantly enriched for any coherent biological processes (Table S2), though this list included individual key regulators of immune function and tolerance such as IFNγ (β = −0.19, adjusted p = 0.02). In contrast, genes upregulated in industrialized environments (n = 635 genes) were enriched for metabolic processes, including pathways related to energy production, nutrient utilization, and biosynthetic regulation (e.g., “small molecule catabolic process" and "carboxylic acid metabolic process", Table S2; Figure 3D). TF enrichment analysis (SI Figure 6; Table S3) identified current industrialized lifestyle associated genes as most strongly enriched for targets of *SPI1* (log2(OR) = 4.52, adjusted p = 1.03×10⁻□), consistent with pathways of immune cell regulation [85] (Table S3). In contrast, non-industrial lifestyle associated genes were enriched for TF targets of *STAT3* and *TP53* (log2(OR) = 3.50, adjusted p = 1.95×10⁻□; log2(OR) = 3.38, adjusted p = 2.38×10⁻□), with roles in cytokine signaling and cellular stress responses [86,87] (SI Figure 6; Table S3). Notably, 31 TFs were shared between the two sets of current lifestyle genes; however, the specific target genes contributing to these enrichments usually differed (SI Figure 6; Table S3).

To identify potential links between lifestyle-associated transcriptional variation and complex traits, we overlapped our differentially expressed gene sets with previous transcriptome-wide association studies (TWAS)—a method that identifies genes where genetic effects on a given trait are mediated through gene expression [88] (SI Figure 7; Table S4). Genes associated with early-life environments showed limited evidence of enrichment, with associations only observed for monocyte count (OR = 1.62, p = 6.75x10^-3^, adjusted p = 0.07, Table S4). Genes associated with current urban environments were not significantly enriched for any complex traits, but genes associated with current non-industrial environments were enriched for autoimmune, asthma, and allergic traits (e.g. diagnosed Hayfever, allergic rhinitis, and eczema: OR = 1.52, adjusted p = 0.016; Table S4). These results are intriguing given previous work linking pathogen- and helminth-rich environments to reduced risk of asthma and allergy, potentially by altering the balance of Th1 cells (which manage intracellular threats to viruses and bacteria) to Th2 cells (which manage extracellular threats from macroparasites and allergens) [26,89–92]. To explore this idea, we asked whether genes up-regulated in non-industrial contexts were enriched within the Th1 and Th2 cell differentiation KEGG pathway (hsa04658), finding that indeed they were (Figure 3C & SI Figure 8, log2(OR) = 2.13, p = 1.39x10^-6^). This enrichment highlights key regulators of T helper cell polarization, for example GATA3 and T-bet, suggesting that current non-industrial environments may influence the balance between pro-inflammatory (Th1) and anti-inflammatory/allergic (Th2) immune responses.

### Current environmental experiences dominate adult gene expression patterns

Given the functional division of gene expression patterns across the life course, we next sought to determine to what degree transcriptomic signatures established in early-life persist despite subsequent environmental change. Because some individuals experienced markedly different environments in early-life versus adulthood (Figure 1C), we focused on individuals at the extremes (upper and lower tertile) of the early-life and current lifestyle distributions, classifying them as “always non-industrial” and subsistence-based (n = 185), “always industrialized” and urban (n = 204), “non-industrial to industrialized" (n = 32), or “industrialized to non-industrial” (n = 36) (Figure 4A). Using gene expression data from the environmentally stable groups (always non-industrial and always industrialized), we trained an elastic net classifier to distinguish these classes of individuals based solely on their transcriptomic profiles (focusing on 5,158 genes not associated with age or ethnolinguistic group at an adjusted p-value of 0.1, AUC=0.94; SI Figure 9). We then applied our classifier to individuals that switched between environments, predicting that, if early-life environments have lasting signatures, then “switching” individuals will be assigned to their early-life group; in contrast, if current experiences determine transcriptional profiles, then switching individuals will be assigned to their current environmental group (Figure 4B).

When we applied our model to individuals who experienced within-lifetime transitions, in either direction, their gene expression profiles were more similar to individuals who shared their current environment rather than their early-life environment (binomial test, p = 6.54x10^-5^, Figure 4C). However, the timing of this transition mattered: for individuals who transitioned from urban to non-industrial as part of “back to roots” movements, we found that time since transition explained heterogeneity in elastic net predictions: specifically, a three category question about time spent in the current community—with options of <1 year, 1-10 years, or >10 years— correlated with how confident the elastic net model was that the focal individual should be classified as non-industrial and subsistence-based (Figure 4D).

### Composite lifestyle score captures broader transcriptomic effects than individual lifestyle components

Our analyses thus far have relied on composite measures of urbanization, market-integration, and industrialization to capture a complex and multifaceted process [93]. Previous work with Orang Asli has shown that these composite measures reflect multiple individual determinants—for example physical activity, consumption of ultraprocessed foods, and helminth infection—that are known to impact inflammation, reactivity, and immune tolerance [41,57,94]. To understand how these proximate determinants contribute to the variation in gene expression we observed in Orang Asli adults, we reran our analyses swapping out our current lifestyle score for six facets found to be associated with this score in previous work and in this sample (SI Figure 10): 1) body roundness index (BRI), a measure that assess body shape and fat distribution [95] (β = 0.505, p = 9.304 x 10^-18^), 2) an index of how much access each individual had to processed, market-derived foods [41] (β = 0.061, p = 2.636 x 10^-13^), 3) an index of how much access each individual had to traditional, forest-derived foods [41] (β = -0.010, p = 2.552 x 10^-49^), 4) objectively measured physical activity levels, operationalized as mean Euclidean Norm Minus One (ENMO) from accelerometry [39] (β = -0.898, p = 1.770 x 10^-2^), 5) eosinophil percentages, the cell type responsible for combating macroparasitic infections (β = -0.0677, p = 4.962 x 10^-2^), and 6) Immunoglobulin E (IgE) levels, an antibody involved in allergic reactions and macroparasitic defenses (β = -0.898, p = 1.770 x 10^-2^) (Table S5; SI Methods). These analyses focused on 660 individuals for whom all of the above measurements were available.

None of the individual factors recapitulated the breadth of gene expression differences captured by the composite lifestyle score, although BRI and ENMO were each associated with a few hundred genes (adjusted p <0.1, n = 229 and n = 159, respectively; Figure 5A). Most genes were specific to either the current lifestyle score, BRI, or ENMO. However, the overlap between genes differentially expressed as a function of lifestyle and BRI (n = 48 genes; p = 3.621x10^-7^), as well as lifestyle and physical activity (n = 7, p = 0.0366), was significantly greater than expected by chance. Even though few individual genes associated with other lifestyle factors passed multiple hypothesis testing correction, effect size estimates were correlated in biologically expected directions (Figure 5B). For example, genes upregulated in more urban, industrialized individuals were also upregulated in individuals with higher BRI (r = 0.26; p < 2.2x10^-16^) and more market-focused diets (r = 0.17; p = 2.75x10^-10^), but downregulated in individuals with higher physical activity levels (r = -0.10; p = 2.48x10^-4^), more traditional diets (r = -0.60; p < 2.2x10^-16^), and putative helminth exposures (higher IgE, r = -0.05; p = 0.079; or higher eosinophil percentages, r = -0.13; p = 3.08x10^-6^). Overall, these results suggest that while individual environmental dimensions (as measured here) explain only a fraction of the transcriptomic variation captured by the composite lifestyle score, they nevertheless likely influence overlapping molecular pathways. These results strongly echo previous work focused on cardiometabolic outcomes [41].

**Figure 5:**
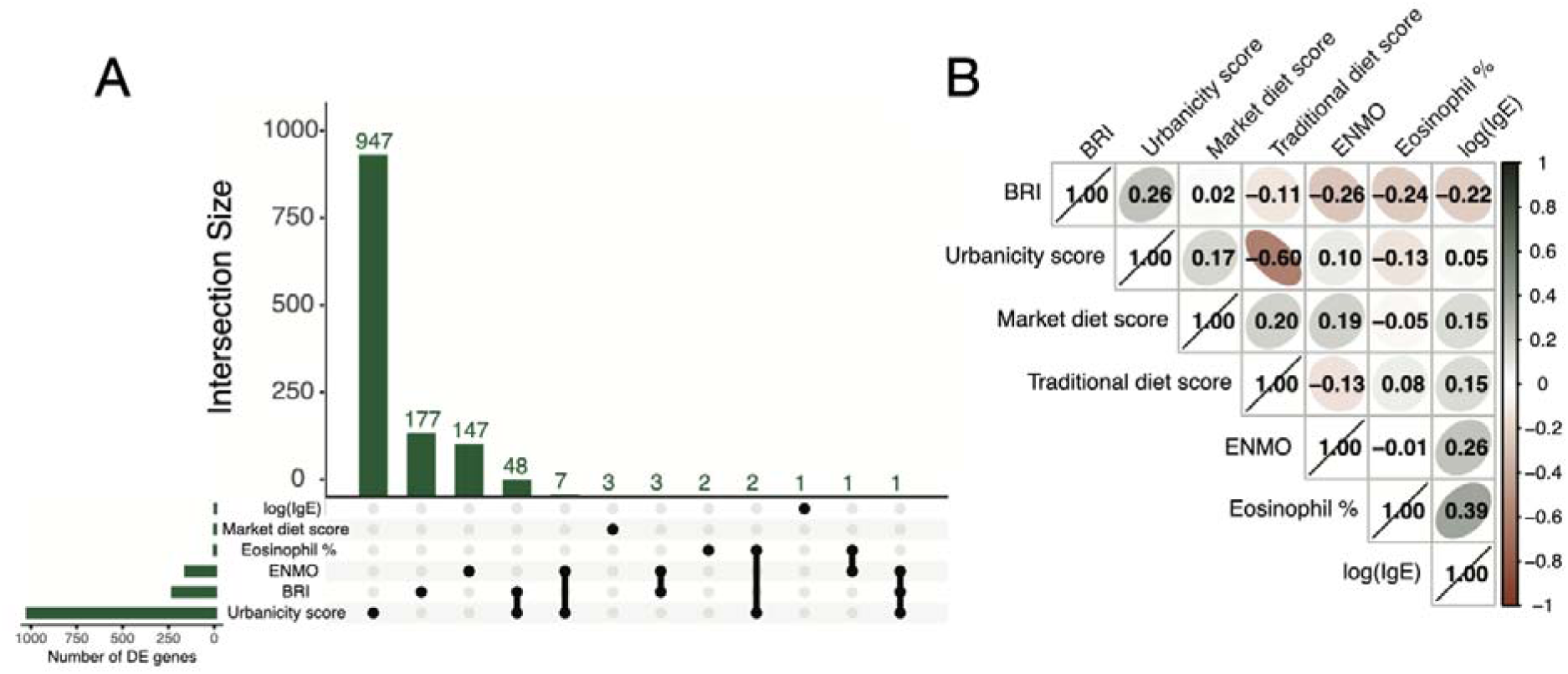
Effects of individual, inflammatory-modulating lifestyle factors on gene expression. (A) Upset plot showing the overlap of genes significantly associated with the current lifestyle score as well as each individual factor at adjusted p < 0.1. (B) Correlation between effect sizes on gene expression and lifestyle factors for any gene significantly associated with at least one lifestyle factor. Abbreviations: ENMO, Euclidean Norm Minus One; BRI, body roundness index; IgE, immunoglobulin E

## Discussion

The human immune system evolved under pathogen pressures, life history trade-offs, and energetic constraints that were substantially different from those experienced in many industrialized environments today. A growing body of work has demonstrated that exposure to urbanized and industrialized conditions across the life course is associated with differences in immune function, yet most studies focus on contemporary environments [19,20,22–25,90,96,97], despite increasing recognition of the developmental origins of health and disease [98,99]. In many cases, this limitation reflects practical constraints on study design: few human populations experience sufficient within-lifetime environmental variation at the time of study to disentangle early-life and current effects. The rapid and heterogeneous lifestyle transitions experienced by the Orang Asli over the past decades presents a valuable opportunity to address this gap. Thus, our study moves beyond asking whether industrialization affects immunity to ask how the timing of environmental change shapes different components of the immune system. Integrating ethnographically informed measures of early-life and current environments with transcriptomic, cellular, and inflammatory phenotypes, our findings support a model in which distinct immune compartments differ in their sensitivity to life course exposures. In general, adaptive immune features were more strongly associated with early-life environments, whereas inflammatory and innate immune phenotypes remained more responsive to current ecological conditions.

Our findings suggest that early-life environments leave durable signatures on adaptive immune traits, particularly T-cell biology (e.g., T cell abundance and IL-8 gene expression and protein levels). This pattern is consistent with experimental work showing that early microbial and antigen exposure shape thymic development and lymphocyte repertoires [30,90], supporting a model in which developmental conditions can establish aspects of adaptive immune architecture that persist into adulthood [26]. This model connects to broader frameworks including developmental origins of disease [99], microbiome-mediated immune priming [27], and the hygiene and “old friends” hypotheses [100–102]. In contrast, current exposure to urban lifestyles was more associated with inflammatory and innate immune system differences, consistent with previous work linking even short-term exposures to proinflammatory factors such as Western diets to similar signatures [25,103,104]. More non-industrial environments promoted immune profiles consistent with ongoing pathogen and macroparasite exposure and greater immunoregulatory activity (especially along Th1 and Th2 pathways), echoing work in populations that experience similar ecologies [105–109]. From a life history perspective, investment in innate versus adaptive immunity may be finely tuned to nutrition, pathogens, and extrinsic mortality risk experienced early in life: in developmental environments where pathogen threats are frequent and potentially deadly, investing heavily in adaptive immune compartments—though costly to maintain, especially in resource constrained environments—is nevertheless likely to provide long-term fitness payoffs [6].

Our comparisons of the relative contributions of early-life and current environments to adult immune phenotypes generally agreed with previous empirical work [110,111], with current lifestyle emerging as the stronger predictor. This observation also agrees with evolutionary theory suggesting that developmental “programming” of adult phenotypes is unlikely to be an adaptive strategy when early life cues are poor or inconsistent predictors of adult conditions [112–115]; the degree to which pre- or post-natal conditions were accurate predictors of adult environments throughout human evolution is unknown, though this is perhaps more likely to have been true for certain facets relevant to immune biology (e.g., pathogen identity or abundance) than others (e.g., resource availability). Relatedly, we were surprised that our analyses to nominate individual lifestyle facets most relevant to immune function did not reveal any single key driver, even though we focused on experimentally well-established predictors [25,103,104,116]. One possible interpretation of these findings is that our composite lifestyle measure may better capture cumulative and coordinated exposures, while for example one week of accelerometry or one-time measures of IgE levels do not fully reflect dynamic exposures. Another non-mutually exclusive explanation is that the immunological consequences of urbanization emerge from the combined effects of many interacting exposures: changes in diet, pathogen exposure, physical activity, sanitation, psychosocial stress, and social structure often occur simultaneously, and their combined effects may better explain immune variation than any one environmental factor considered in isolation.

Despite identifying generally strong effects of current lifestyle, our elastic net results suggest that immune signatures do not respond instantaneously to environmental transitions: people who transitioned from urban, industrialized contexts to non-industrial lifeways retained time-dependent signatures of prior exposures. This temporal persistence is consistent with concepts such as trained immunity—in which innate immune cells can acquire epigenomic reprogramming that allows them to mount a heightened, non-specific response to secondary stimuli [117,118]. In other words, while current conditions will generally reshape immune function, there are putative mechanistic explanations for some amount of short-term memory of recent environments.

Our study has several limitations. First, although observational studies such as ours remain essential for understanding real-world lifestyle transitions, establishing causality remains challenging. A growing body of experimental work has begun to bridge this gap by demonstrating casual effects of environmentally derived exposures on immune function in animal models and *ex vivo* systems [22,90]. For example, Stein and colleagues treated mice with dust collected from Amish (traditional farmers) and Hutterite (industrialized farmers) households, to show causal effects of these exposures on immune function that echoed differences in inflammation, asthma, allergy observed between Amish and Hutterite children [90]. Similar approaches will be valuable for identifying the mechanisms underlying the associations observed here. Second, immune phenotypes were measured only in adulthood, and our lifestyle surveys captured broad conditions in “childhood” and “currently” rather than pinpointing the timing of specific exposures. Consequently, we cannot determine precisely when or which environmental factors, including pathogens, diet, psychosocial stress, or other correlated exposures, drive the observed associations, leaving clear directions for future work. In particular, longitudinal studies incorporating detailed exposure assessment and immune system monitoring will be important for resolving these questions. Finally, given the distinct results we observed for adaptive versus innate immune compartments, future work using single-cell RNA sequencing will be valuable for resolving lifestyle-associated effects within individual immune cell types.

Overall, our findings suggest that different components of the immune system encode and retain environmental experience over different timescales. Consequently, immune variation cannot be understood solely in terms of present-day environments. This perspective perhaps helps reconcile why individuals exposed to similar industrialized or other types of environments in adulthood can nevertheless exhibit markedly different health conditions and disease risks. The combination of developmental embedding and adult plasticity we observe is relevant for explaining why industrialized populations exhibit elevated risk of inflammatory and metabolic diseases, including cardiovascular disease, type 2 diabetes, and autoimmune disorders, but critically suggests that a path toward protection could include a focus on current as well as early life conditions [9–16]. From a policy and community perspective, the substantial influence of current environments suggests that most of the health consequences of industrialization remain physiologically reversible and poised for intervention. For the Orang Asli in particular, this observed plasticity provides molecular support for the idea that rejection of urban, industrialized lifeways and transitions “back-to-roots” can produce measurable physiological and health benefits.

## Materials and Methods

### Study population and lifestyle variation

The Indigenous Orang Asli of Peninsular Malaysia comprise multiple ethnolinguistic groups that vary in their degree of urbanization, market integration, and subsistence practices [37,119]. Over recent decades, rapid socioeconomic development in Malaysia has produced marked heterogeneity in environmental exposures across Orang Asli communities, who now range from remote rainforest villages that continue to rely heavily on hunting, gathering, horticulture, and fishing to communities integrated into industrialized and urban environments [120–124]. Participants spanning this lifestyle gradient were enrolled through the Orang Asli Health and Lifeways Project (OA HeLP), an ongoing collaboration between international scientists, local health care and Indigenous rights organizations, and communities to understand lifestyle effects on health and well-being. Data used in this study were collected between June 2022 and December 2024 from adults (≥18 y old). During this time, researchers visited Orang Asli villages throughout Peninsular Malaysia. At each location, informed consent was collected at multiple levels: first by describing the project to the community as a whole and seeking the permission of community leaders, and subsequently through individual review of the protocol followed by formal, written consent. Data collection involved surveys about both early-life and current environments, especially as these environments relate to lifestyle, acculturation, market-integration, and urbanization (see SI Materials and Methods, as well as [37]).

Early-life environmental variation was quantified using retrospective questionnaire data covering parental subsistence activities, household assets, housing materials, and dietary practices. Principal components analysis (PCA) was used to summarize these variables, and the first principal component was retained as an index of early-life lifestyle [15]. Current lifestyle was quantified using a previously validated location-level score based on infrastructure access, population density, household assets, education, and subsistence activities [16,57]. Details regarding questionnaire content and processing, and derivation of lifestyle scores, are provided in SI Materials and Methods.

### Anthropometric, dietary, and immunological measurements

As part of ongoing OA HeLP mobile clinics, anthropometric and clinical measurements were collected during medical visits using standardized protocols [37]. Body roundness index (BRI) was calculated from waist circumference and standing height as a continuous estimate of body shape and adiposity [95,125]. Dietary information was collected using structured food-frequency questionnaires assessing both traditional and market-based dietary practices. Separate indices representing market-based and traditional dietary consumption were derived from scaled intake frequencies (see SI Materials and Methods). Physical activity was quantified using wrist-worn accelerometers worn continuously for approximately one week [39].

Peripheral blood samples were collected from participants for hematological, immunological, and transcriptomic analyses. Whole blood samples were used to assess white blood cell composition via automated hematology analyzers (HemoCue) when available or manual differential counts from a blood smear otherwise. Whole blood was also drawn into CPT tubes for PBMC and plasma isolation. Plasma IgE concentrations were quantified by ELISA (DRG International). Plasma cytokines and inflammatory biomarkers were measured by the Duke Biomarkers Core at the Duke Molecular Physiology Institute using multiplex electrochemiluminescent immunoassays (Meso Scale Discovery platform). Detailed assay procedures and processing steps are described in SI Materials and Methods.

### Bulk and single-cell RNA sequencing

Total RNA was extracted from peripheral blood mononuclear cells (PBMCs) and used for poly(A)-selected library preparation (using kits from Zymo research and New England Biosciences, respectively). Prepared libraries were paired-end sequenced on the Illumina NovaSeq X platform. Reads were aligned to the human reference genome (hg38) [126], quantified at the gene level [127], filtered to remove low-quality samples and lowly expressed genes, and normalized using the voomWithQualityWeights function from the R package limma [128]. Our processing and filtering steps resulted in a final dataset of 922 individuals and 9,993 expressed genes for analysis. Additional quality-control procedures and processing details are provided in SI Materials and Methods.

To generate a reference atlas for immune cell deconvolution, PBMCs from four individuals were profiled using the 10x Genomics Chromium Flex platform. Single-cell transcriptomic data were processed using Cell Ranger and Scanpy, with dimensionality reduction, clustering, and reference-based cell type annotation based on standard workflows [129,130]. High-confidence cell type signatures derived from the single-cell data were used for reference-based deconvolution of bulk RNA-seq profiles [131].

### Statistical analyses

Associations between lifestyle variables and physiological, immunological, and transcriptomic outcomes were assessed using linear regression and linear mixed-effects models implemented in R version 4.4.2. For analyses of immune cell proportions and circulating biomarkers, current lifestyle score and early-life lifestyle score were included as predictors of interest while controlling for age, sex, and assay batch. Multiple-testing correction was performed using the Benjamini–Hochberg false discovery rate (FDR) [132], with FDR adjusted p < 0.1 considered significant. Cytokine concentrations were log -transformed and standardized to mean 0 and unit variance prior to analysis. Normalized expression values were obtained using voom with sample-specific quality weights, and technical effects (sequencing batch, uniquely mapped reads, monocyte percentage, and lymphocyte percentage) were regressed out using limma [128]. Residualized gene expression was then modeled for each gene using linear mixed effects models implemented in EMMREML [133]. Models included current lifestyle score, early-life lifestyle score, age, sex, and self-identified ancestry group as fixed effects, while accounting for genetic relatedness among individuals using a genetic relatedness matrix derived from RNA-seq genotypes [134]. We applied FDR correction across all genes and considered an FDR adjusted p < 0.1 to be significant.

Functional enrichment analyses were performed on lifestyle-associated genes using Gene Ontology annotations in clusterProfiler [135]. Transcription factor enrichment was evaluated using curated TF–target interactions from the TRRUST v2 database and Fisher’s exact tests [82]. We additionally tested whether lifestyle-associated genes were enriched for genes in the KEGG Th1 and Th2 cell differentiation pathway (hsa04658) using Fisher’s exact tests [136]. To assess potential links with human disease, we tested for enrichment of genes implicated in published probabilistic transcriptome-wide association studies (PTWAS) of noncommunicable disease traits using Fisher’s exact tests [88]. Finally, to evaluate whether transcriptomic profiles could predict lifestyle categories, elastic net logistic regression models were trained using gene expression data after excluding genes associated with age or ancestry (FDR adjusted p < 0.1, leaving 5,158 genes). Models were trained using leave-one-out cross-validation [137,138]. Additional details regarding preprocessing, model specification, deconvolution procedures, enrichment analyses, and classifier training are provided in SI Materials and Methods.

### Ethics

Procedures for this study have been reviewed and approved by the Medical Review and Ethics Committee of the Malaysian Ministry of Health (protocol ID: NMRR-20-2214-55565), the Malaysian Ministry of Economy (permit ID: EPU 40/200/19/3911), the Malaysian Department of Orang Asli Development (permit ID: JAKOA.PP.30.052 JLD 21 (98)) and the Institutional Review Boards of the University of New Mexico (protocol ID: 14420) and Vanderbilt University (protocol ID: 212175). Throughout the project, we have followed established principles for ethical biomedical research among Indigenous communities, including fostering collaboration, building cultural competency, being transparent about research practices, supporting capacity building and disseminating findings [139].

## Acknowledgements

We thank Orang Asli participants and communities for their interest, support, hospitality and willingness to work with us on this project. We thank the OA HeLP field team and volunteers, whose expertise and commitment made this work possible. We are grateful to the local organizations and institutions that we work with, namely the Center for Orang Asli Concerns, the Malaysian Red Crescent Society, Federation of Private Medical Practitioners’ Associations of Malaysia (Drs4All program), Hospital Orang Asli Gombak, and Universiti Malaya. This work was supported by the National Science Foundation (BCS-2142090), the Canadian Institute for Advanced Research (Azrieli Global Scholars Program), the Pew Charitable Trusts (Pew Biomedical Scholars Program), and the Vanderbilt University Evolutionary Studies Initiative (NIH/NIGMS 1T32GM150407). The funders had no role in study design, data collection and analysis, decision to publish or preparation of the manuscript.

## Data and Code Accessibility

OA HeLP prioritizes minimizing risk to study participants and follows the ‘CARE Principles for Indigenous Data Governance’ and the FAIR Guiding Principles for scientific data (emphasizing Findability, Accessibility, Interoperability and Reusability). Individual-level data are stored on Zenodo in a protected repository and available through restricted access (https://zenodo.org/records/21535682; DOI 10.5281/zenodo.21535681). Requests for de-identified data must include a detailed application and procedures for data security, privacy and minimizing potential harm. A template for data use agreements is provided at https://lea-lab.org/resources.html. All code for analyses presented here are available on GitHub (https://github.com/laylabrassington/ELvsCL); individual scripts associated with each analysis are described in the supplementary materials associated with the manuscript.

