## Supplementary Materials for "Early and current environments exert distinct effects on immune function in the Orang Asli"

**This file includes:**

- Supplementary Materials and Methods
- SI Figure 1: Age and sex distribution of participants.
- SI Figure 2: Estimation of blood cell-type composition in Orang Asli individuals using an Orang Asli-specific single-cell RNA-seq reference panel.
- SI Figure 3: Comparison of cell-type deconvolution results generated using a non-Orang Asli versus an Orang Asli-specific single-cell reference panel.
- SI Figure 4: Sharing and magnitude of early-life and current lifestyle effects on adult gene expression.
- Figure S5: Permutation analysis of early-life (EL) and current lifestyle (CL) gene expression model results.
- Figure S6: Transcription factor target enrichment among lifestyle-associated gene sets.
- Figure S7. Overlap between lifestyle-associated gene sets and transcriptome-wide association study (TWAS) hits.
- Figure S8: Overlap between current subsistence-level genes and the T helper cell differentiation pathway.
- Figure S9: Predictive performance of lifestyle classification models under alternative gene filtering criteria.
- Figure S10: Associations between current lifestyle score and four aspects of lifestyle relevant to immune function.
- Supplementary References

**The following tables are provided in a separate file:**

- Table S1: Variables included in lifestyle score construction
- Table S2: Results from gene ontology (GO) analyses for different gene sets; the top 20 results are reported for each analysis
- Table S3: Results from transcription factor binding enrichment analyses for different gene sets
- Table S4: Results from TWAS enrichment analyses for different gene sets
- Table S5: Results from linear models testing for associations between the current lifestyle index and individual lifestyle facets
- Table S6: Bulk mRNA-seq metadata and metrics
- Table S7: Single cell mRNA-seq metadata and metrics
- Table S8: Top 10 marker genes per cell type used for deconvolution of bulk mRNA-seq data

Supplementary Materials and Methods

**The Orang Asli of Peninsular Malaysia**

The Indigenous Orang Asli comprise <1% of the population of Malaysia (totaling ∼238,000 people) and include at least 18 distinct ethnolinguistic groups, which are typically divided into three broad categories: the Semang/Negrito (traditionally nomadic hunter-gatherers speaking northern Aslian languages), Senoi (traditionally horticulturalists speaking central Aslian languages), and Proto-Malay (traditionally practitioners of mixed subsistence economies speaking Austronesian language dialects) [[2,3]](https://paperpile.com/c/zOgmfo/q553+jFcU). In addition, each of the 18 different ethnolinguistic groups have rich and varied histories that have resulted in distinct religions, coresidence patterns, marriage, practice, mobility patterns, and other aspects of socioecology. Although subtle genetic variation exists among ethnolinguistic groups, Orang Asli populations are relatively genetically similar compared with surrounding Asian populations [[4]](https://paperpile.com/c/zOgmfo/PNwu).

Over the last half-century, Orang Asli communities have experienced substantial lifestyle changes associated with rapid socioeconomic development in Malaysia. First, expansion of industries focused on plantation agriculture (particularly oil palm and rubber) and natural resource extraction (particularly timber, tin, and petroleum) since the 1970s and 1980s, when the interior regions of Peninsular Malaysia became more accessible [[5–7]](https://paperpile.com/c/zOgmfo/cxxGW+BxEk+iCLgY), have led to both significant deforestation and increased infrastructure [[8–12]](https://paperpile.com/c/zOgmfo/uekq+fyqO+copW+KXYD+U6Jz). Orang Asli have remained relatively marginalized throughout this process, which has prompted an increased dependence on wage labor and the cash economy among many individuals as traditional lands are threatened [[13,14]](https://paperpile.com/c/zOgmfo/YMdy+hySd). Second, efforts by the Malaysian government to assimilate the Orang Asli into mainstream culture [[14–16]](https://paperpile.com/c/zOgmfo/o2Mh6+Dnk7+hySd), including through development and resettlement schemes focused on “modern” facilities [[13,14]](https://paperpile.com/c/zOgmfo/YMdy+hySd), have created shifts toward more urban, acculturated environments in certain geographic areas [[17–20]](https://paperpile.com/c/zOgmfo/eqTWE+5eL1U+9jqCT+VM2D). Importantly, due to the pace of these changes, variation exists in how long people have engaged in wage labor or lived in urban environments, with some people having made these transitions only recently and others having been exposed to such contexts for their entire lives. Third, some ethnolinguistic groups and individual villages have resisted sedentarization, interaction with the cash economy, and the authority of industry, government, and other entities, with some communities rejecting these lifestyles outright and moving from government settlements “back to roots” (i.e., returning to remote, rainforest villages) [[13,21]](https://paperpile.com/c/zOgmfo/YMdy+3gugB). As a result of these trends, there are currently remote communities located in the rainforest that rely heavily on the availability of natural resources [[22,23]](https://paperpile.com/c/zOgmfo/gNO91+G8MwH), as well as communities now immersed in fully industrialized economies and urban environments. Further, variation exists in how long people have lived in each environment, with some people having made these transitions only recently and others having been exposed to such contexts for their entire lives.

**Data collection overview and lifestyle score construction**

The Orang Asli Health and Lifeways Project (OA HeLP) is an interdisciplinary collaboration focused on understanding how lifestyle change is influencing health and well-being among Orang Asli. The project is composed of an international research team of anthropologists, biologists, and physicians. A detailed protocol for OA HeLP is provided elsewhere [[24]](https://paperpile.com/c/zOgmfo/sFaE), but data collection includes diverse information from questionnaires, anthropometry, health assessments, and biospecimens. The data used for this study were collected between June 2022 and December 2024 from Orang Asli villages throughout Peninsular Malaysia. At each location, informed consent was collected at multiple levels: first by describing the project to the community as a whole and seeking the permission of community leaders, and subsequently through individual review of the study procedures followed by formal, written consent. For this study, we worked with communities spanning the following ethnolinguistic groups: Batek, Batek Nong, Jahut, Jakun, Jehai, Kensiu, Kintaq, Lanoh, Mah Meri, Semaq Beri, Semai, Semelai, Temiar, and Temuan. All participants completed questionnaires assessing both early-life (centered broadly around childhood experiences) and current environments, including dietary practices, housing materials, parental occupations, household assets, subsistence activities, and indicators of market integration and urbanization (the full list of relevant variables is provided in Table S1) [[24]](https://paperpile.com/c/zOgmfo/sFaE).

Early-life lifestyle score: To create an index of early life lifestyle, we applied a dimension reduction approach to our early life questionnaire data (Table S1) as previously proposed in the literature [[25]](https://paperpile.com/c/zOgmfo/DpYa) and validated for Orang Asli [[26]](https://paperpile.com/c/zOgmfo/4e2k). Specifically, unordered categorical responses were converted into binary indicator variables representing the presence or absence of each response category. Ordered categorical responses were converted into numeric scales. For questionnaire items concerning parental occupation during childhood, missing values resulting from the death or absence of one parent were replaced with values from the other parent (n = 76 participants). Additional missing data affecting one or two questionnaire items among 25 participants were imputed using the R package mice with LASSO-based imputation [[27]](https://paperpile.com/c/zOgmfo/9fUo). The resulting dataset was summarized using principal components analysis implemented with the prcomp function from the stats package in R. The first principal component explained 21% of total variation and was retained as a summary measure of early-life environmental variation based on scree plot inspection and interpretability of variable loadings. See [[26]](https://paperpile.com/c/zOgmfo/4e2k) for additional details.

Current lifestyle score: To create an index of current lifestyle, we applied a location-based approach as previously proposed in the literature [[28]](https://paperpile.com/c/zOgmfo/8QFw) and validated for Orang Asli [[1]](https://paperpile.com/c/zOgmfo/vC7r). The score included measures of population density; the proportion of households with access to electricity and sewage systems; the proportion of households owning mobile phones and televisions; the proportion of individuals engaged in subsistence practices; infrastructure available to access the village; and the proportion of individuals with some formal education among those aged >40 years and <40 years. Individual- and household-level variables were aggregated within each community to generate village-level scores, with higher scores reflecting greater industrialization. The score is an additive measure which includes: weighted population density in a 5km grid (across a 10 point scale), 10 points- 10*(proportion of population involved in non-wage labor), 5 points* proportion of households with flush toilets, 5 points* proportion households with electricity, 5 points* proportion households with television, 5 points* proportion households with mobile phone, 10*proportion >40yo with some education, and 10*proportion <=40yo with some education. See Table S1 for the list of components and [[1,28]](https://paperpile.com/c/zOgmfo/vC7r+8QFw) for additional details.

Market diet index: This index was created to determine the extent to which individuals consume a market derived diet. The index consists of frequency of sugar, salt, and cooking oil consumption as assessed from our current life questionnaire data. It is coded such that higher values indicate more consumption of market-derived foods, and it has previously been validated for Orang Asli [[1]](https://paperpile.com/c/zOgmfo/vC7r).

Traditional diet index: This index was created to determine the extent of traditional foods consumed. The index specifically consists of frequency of rice, manioc, wild meat, and wild fish consumption. It is coded such that higher values indicate more consumption of traditional foods, and has been previously validated for Orang Asli [[1]](https://paperpile.com/c/zOgmfo/vC7r).

**Accelerometry data collection and preprocessing**

At the conclusion of mobile health clinics, participants were invited to wear accelerometers (Axivity AX3) on the non-dominant wrist for approximately one week. Devices were configured to record acceleration data at 100 Hz with an 8 g dynamic range. Accelerometry data collection occurred outside of mobile health clinic periods to minimize behavioral changes associated with the presence of researchers. Raw accelerometry data were downloaded and preprocessed to remove days recorded before or after the wear period. Data were processed using the GGIR package (v3.2.6) in R [[29,30]](https://paperpile.com/c/zOgmfo/dam7+dzwQ) with default autocalibration and imputation procedures. Non-wear time was estimated using built-in GGIR algorithms, and valid days were defined as those with ≥16 h of detected wear time. Physical activity was quantified using the mean Euclidean Norm Minus One (ENMO) averaged across 24-h periods and expressed in milligravity units (mg). Participant-level activity estimates were calculated by averaging ENMO values across all valid days. See [[31]](https://paperpile.com/c/zOgmfo/Mtex) for additional details.

**Blood sample collection and measures of immune variation**

For all participants in this study, approximately 8 mL of venous blood was collected into CPT tubes for PBMC isolation and downstream bulk mRNA-seq following manufacturer protocols. PBMCs were preserved in DNA/RNA Shield (Zymo Research) and plasma was isolated following separation; all samples were immediately stored in liquid nitrogen followed by long-term storage at −80 °C. During mobile clinics, whole blood was also used to conduct a five-part white blood cell differential using an automated hematology analyzer (HemoCue WBC DIFF System), with manual differential counts from blood smears used for a small subset of participants for whom this assay was unavailable. For downstream data analysis, eosinophil percentages were log₂-transformed after addition of a pseudocount of 0.1 to reduce skewness.

For a small subset of participants, 4-5 mL of whole blood was set aside for PBMC cryopreservation for single cell mRNA-seq, with less sample devoted to fixation in DNA/RNA Shield as described above. Specifically, peripheral blood mononuclear cells were isolated from fresh whole blood using density gradient centrifugation with SepMate™ tubes (STEMCELL Technologies) according to the manufacturer’s protocol. Briefly, 4-5 mL of whole blood collected in EDTA tubes was diluted 1:1 with PBS containing 2% FBS. The diluted blood was layered over 3.5 mL of density gradient medium in SepMate tubes and centrifuged at 1,200 x g for 10 min at room temperature with the brake on. The plasma layer was carefully removed, leaving approximately 1-2 mL above the PBMC interface. The PBMC layer was transferred to a fresh 15 mL conical tube, washed with PBS + 2% FBS, and pelleted by centrifugation at 250-300 x g for 10 min. For cryopreservation, cell pellets were gently resuspended in chilled FBS, and an equal volume of 2x freezing medium (20% DMSO in FBS) was added dropwise while swirling to a final concentration of 10% DMSO. Approximately 0.75 mL of the resulting suspension was dispensed into each of two prechilled cryovials. Cryovials were placed in a controlled-rate freezing container that was put in a liquid nitrogen tank for 24 hours, then taken out and subsequently stored in liquid nitrogen for long-term preservation.

For a subset of plasma samples collected from CPT tubes, plasma IgE concentrations were measured using enzyme-linked immunosorbent assays (DRG International kit EIA-1788). IgE values were log₂-transformed prior to analysis. For an additional subset of plasma samples, immune cytokine and biomarker concentrations were measured by the Duke Biomarkers Core at the Duke Molecular Physiology Institute using multiplex electrochemiluminescent immunoassays (Meso Scale Discovery platform). Thirteen inflammatory and metabolic biomarkers were quantified: VPLEX C-Reactive Protein (CRP) (Cat K151STD), VPLEX Leptin (Cat K151V5D), RPLEX Lipopolysaccharide-binding protein (LBP) (Cat K151K5R), and VPLEX Proinflammatory Panel 1:, IFN-γ, IL-1β, IL-2, IL-4, IL-6, IL-10, IL-12, and IL-13, TNF-α (Cat K15049D). See additional curation details in ”Immune biomarker statistical analyses”. The mean intra- and inter-assay coefficients of variation for all measured biomarkers were <15%.

**Bulk and single cell mRNA sequencing**

Bulk mRNA-seq: Total RNA was extracted from peripheral blood mononuclear cells (PBMCs) using either the Zymo Quick-DNA/RNA Miniprep Kit or the Zymo Quick-RNA 96 Kit (Zymo Research), following the manufacturer’s instructions. RNA libraries were generated using the NEBNext Ultra II RNA Library Prep Kit (New England Biolabs) in conjunction with the mRNA Magnetic Isolation Module, following the manufacturer’s protocol with an input of 150 ng of total RNA per sample. Libraries were pooled in equimolar concentrations and sequenced on the Illumina NovaSeq X Series platform using paired-end 150 bp reads at either the Vanderbilt VANTAGE Genomics Core or The Translational Genomics Research Institute (TGen).

Single cell mRNA-seq: Cryopreserved PBMCs from four individuals were used for single cell mRNA-seq. All four samples were from female participants with ages ranging from 19-24.4 at the time of sample collection. The individuals were from four different sampling locations representing four ethnolinguistic groups. Cryopreserved PBMCs were thawed rapidly at 37C, washed in RPMI 1640 medium supplemented with 10% FBS, and filtered to remove debris prior to loading. Single-cell suspensions were processed using the 10x Genomics Chromium Flex Gene Expression GEM-X platform (4 x 4 format) following the manufacturer’s instructions. Libraries were constructed using the 10x Chromium Flex Gene Expression kit, which employs microfluidic encapsulation of individual cells with barcoded gel beads for reverse transcription and library generation. Libraries were sequenced on the Illumina NovaSeq X series platform using paired-end 150 bp reads on 10B flow cells, targeting a depth of approximately 20,000 reads per cell. Library prep and sequencing was performed at the Vanderbilt VANTAGE Core facility.

**Bulk mRNA-seq data processing**

Associated script: https://github.com/laylabrassington/ELvsCL

Adapter and low-quality base trimming were performed with Cutadapt (v1.16) using a minimum read length of 20 bp and a quality cutoff of 20 [[32]](https://paperpile.com/c/zOgmfo/KEtB). Trimmed reads were aligned to the human reference genome (hg38) using STAR (v2.5.4b) with default parameters and a maximum multimapping threshold of one read per locus [[33]](https://paperpile.com/c/zOgmfo/2KwS). Gene-level counts were quantified using HTSeq-count (v0.9.1) in intersection-nonempty mode with reference annotations from NCBI RefSeq [[34]](https://paperpile.com/c/zOgmfo/AYfC). Resulting count matrices were merged across sequencing batches and filtered to remove low-quality libraries (total reads <1 million, <70% uniquely mapped reads, or >10% hemoglobin contamination). Only protein-coding genes were retained, and hemoglobin transcripts (HBA1, HBA2, HBB) were excluded. Transcripts per million (TPM) values were calculated using gene lengths derived from GTF annotations, and low-abundance genes (median TPM < 2) were removed, leaving 9,993 genes for analysis. The sex of each sample was verified using expression of Y chromosome genes identified from Ensembl BioMart annotations [[35]](https://paperpile.com/c/zOgmfo/0mPk). Specifically, the proportion of reads mapping to protein-coding genes located on chromosome Y was calculated relative to total protein-coding gene counts for each sample, and inferred sex was compared with reported biological sex to confirm metadata concordance. Normalized expression values were obtained using voom with sample-specific quality weights (voomWithQualityWeights) [[36]](https://paperpile.com/c/zOgmfo/YWri), and technical effects (sequencing batch, uniquely mapped reads, monocyte percentage, and lymphocyte percentage) were regressed out using limma (v3.62.2) [[36]](https://paperpile.com/c/zOgmfo/YWri). The resulting quality-weighted normalized, residualized expression matrix was used for downstream differential expression analyses; for deconvolution analysis, we used a version of the matrix in which we did not regress out cell percentages. See Table S6 for additional per sample metadata.

**Genotyping of bulk mRNA-seq reads and genetic relatedness matrix generation**

Associated script: https://github.com/laylabrassington/rnaseq_genotyping

To generate sample-level genotype calls from bulk mRNAseq reads, aligned BAM files were processed using a standardized variant-calling pipeline based on GATK (v4.1.4.0) [[37]](https://paperpile.com/c/zOgmfo/0UVW) and SAMtools (v1.9) [[38]](https://paperpile.com/c/zOgmfo/OUs8). Briefly, coordinate-sorted BAMs were filtered to retain only autosomal reads and indexed with SAMtools. Duplicate reads were identified and marked using Picard (v2.18.27) [[37]](https://paperpile.com/c/zOgmfo/0UVW), and read group information was added to ensure compatibility with downstream GATK tools. SplitNCigarReads was used to adjust spliced alignments for RNAseq data. Base quality score recalibration (BQSR) was performed using known high-confidence SNPs from the 1000 Genomes Project phase 1 as reference sites. Variant calling was conducted per sample using GATK HaplotypeCaller in gVCF mode, followed by variant quality filtering to remove calls with Fisher Strand (FS > 30.0) or low quality-by-depth (QD < 2.0). The resulting per-sample filtered VCFs were zipped, indexed, and merged across all individuals with bcftools (v1.18). Only variants with the PASS filter flag were retained for subsequent analyses. The merged multi-sample VCF was further filtered to remove variants with excessive missingness (allele number < 50% of samples), minor allele frequency (MAF) < 1%, or Hardy-Weinberg equilibrium (p<1×10−6), using PLINK (v1.9). After all filtering steps, high-quality biallelic SNPs were retained for genotype-based quality control. Principal component analysis (PCA) was performed using PLINK to quantify ancestry and identify potential outliers [[39]](https://paperpile.com/c/zOgmfo/41zC). Pairwise relatedness and kinship coefficients were computed using KING (v2.3.2) to confirm sample independence and identify potential duplicates [[40]](https://paperpile.com/c/zOgmfo/hTHm). The final high-confidence genotype matrix and kinship estimates were used to generate a kinship matrix for inclusion in linear mixed effects models in downstream analysis.

**Single-cell mRNA sequencing and processing**

Associated script: https://github.com/laylabrassington/scDeBulkR

Raw sequencing data were processed using the 10x Genomics Cell Ranger pipeline to generate gene-barcode count matrices. Subsequent analysis was performed in Python using Scanpy (v1.x) [[41]](https://paperpile.com/c/zOgmfo/QEfB). Individual samples were imported and merged after alignment of metadata. Low-quality cells were removed based on multiple quality-control metrics: cells with >5% mitochondrial gene expression, >2% hemoglobin (HBB) transcripts, <300 detected genes, or >1% of counts derived from cell cycle-associated genes were excluded. Potential doublets were identified and filtered using Scrublet (doublet score ≥0.2) [[42]](https://paperpile.com/c/zOgmfo/84jV). Mitochondrial genes were excluded from the dataset, and only genes expressed in >1% of cells in at least one sample were retained. Counts were normalized to 10,000 per cell, log-transformed, and scaled to unit variance. Principal component analysis (PCA) was used for dimensionality reduction, and batch correction was performed using Harmony based on sample identity [[43]](https://paperpile.com/c/zOgmfo/OMGY). The resulting corrected PCs were used to compute a neighborhood graph, followed by Leiden clustering (resolution = 1) and UMAP visualization. Cell type annotation was performed using reference-based prediction with CellTypist (Immune_All_Low and Immune_All_High models) [[44]](https://paperpile.com/c/zOgmfo/EU1T). Cells with a confidence of less than 0.7 were removed. See Table S7 for additional per sample information.

**Bulk mRNA-seq deconvolution and statistical analysis:**

Associated script: https://github.com/laylabrassington/scDeBulkR

Reference-based deconvolution of bulk mRNA-seq data was performed using single-cell-derived transcriptional signatures as input. Normalized, quality-weighted expression matrices were obtained from the limma-voom pipeline and imported into R as an ExpressionSet object. Single-cell RNA-seq data were filtered to include only high-confidence cells (classification probability ≥0.7) and further refined by excluding discordant k-nearest neighbor (kNN) assignments and outliers identified by DBSCAN clustering. Additionally, cell type identity was further confirmed by evaluating the top differentially expressed genes per cluster using Scanpy’s “FindAllMarkers” and filtering cells based on expression level of DEG genes. Dendritic cells were excluded from deconvolution and downstream analyses due to low counts. Marker genes for each annotated cell type were identified using Scanpy’s rank_genes_groups function (method = t-test), and only genes shared between the single-cell and bulk datasets were retained. Cell-type-specific marker sets were generated using adjusted p-value and specificity cutoffs of 0.1 (Table S8). These signatures were aggregated into reference matrices using Seurat’s AggregateExpression function [[41]](https://paperpile.com/c/zOgmfo/QEfB). Bulk RNA-seq samples were deconvolved against the single-cell reference using CIBERSORTx within bseqsc [[45,46]](https://paperpile.com/c/zOgmfo/K6ek+TFMf). The bseqsc implementation resulted in cell-type proportion matrices which were merged and saved as combined R objects for downstream correlation and validation analyses.

Deconvolution outputs were validated by (i) comparing inferred immune cell proportions against matched complete blood count (CBC) data and (ii) benchmarking against publicly available 10x Genomics PBMC datasets (see <https://www.10xgenomics.com/datasets>) to confirm biological consistency and cell-type specificity (Figure S3). The reference panel derived from a non-Orang Asli population yielded similar correlation strengths with cell type proportions derived from whole blood (lymphocytes: r = 0.35, p = 1.52×10^-28^; monocytes: r = 0.08, p = 0.02). However, the non-matched panel produced inflated estimates, with some estimated cell type proportions falling outside expected biological ranges (e.g., lymphocytes up to ~93% vs ~78% with the population-matched panel). Together, these results highlight that while non-matched reference panels may preserve rank-order relationships, population-matched references are critical for generating biologically realistic absolute estimates (Figure S3).

To investigate how current and early-life lifestyle are associated with peripheral immune composition, we modeled deconvoluted immune cell proportions across individuals as a function of lifestyle and demographic covariates. For each participant, the relative abundance of 6 major immune cell populations was derived from single-cell transcriptomic data using the supervised deconvolution framework described above. We fit linear models for each immune cell type controlling for age and sex. Both current lifestyle score (scaled continuous score) and early-life lifestyle score (first principal component of early environment variables) were entered simultaneously to estimate their independent contributions. For each cell type, regression coefficients and p-values were extracted. To account for multiple testing across all cell types and predictors, false discovery rate (FDR) correction was applied using the Benjamini-Hochberg method [[47]](https://paperpile.com/c/zOgmfo/xwSh). Associations with FDR-adjusted p < 0.1 were considered significant. The direction of each effect was interpreted according to the sign of the regression coefficient (β): positive β values indicated higher cell type proportions in more urbanized environments, and negative β values indicated higher proportions in less urbanized environments.

**Immune biomarker statistical analyses**

Associated script: https://github.com/laylabrassington/ELvsCL/blob/main/scripts/modeling.Rmd

Plasma cytokine and biomarker data were analyzed to test associations with current and early-life lifestyle measures. For 3 individuals with repeated samples, cytokine concentrations were averaged across replicates to obtain a single mean value per individual. Values below the LLOD were removed and outlier values were removed via visual inspection of the distribution of the data. All cytokine concentration values were log₂-transformed. Transformed variables were then standardized (z-score scaling) to enable comparison of effect sizes across cytokines. Associations between circulating cytokine levels and environmental variables were modeled using multiple linear regression. For each cytokine, each model was fitted using the base R linear model function testing for the effect of early-life and current lifestyle controlling for age, sex, and batch (assay plate). A Benjamini-Hochberg false discovery rate (FDR) correction [[47]](https://paperpile.com/c/zOgmfo/xwSh) was applied across all cytokine-predictor associations and associations were considered significant at FDR < 0.1.

**Differential expression analyses**

Associated script: https://github.com/laylabrassington/ELvsCL/blob/main/scripts/modeling.Rmd

Main analysis: Gene expression was analyzed using linear mixed models implemented with the EMMREML R package to account for genetic relatedness among individuals [[48]](https://paperpile.com/c/zOgmfo/EZbl). Models included current lifestyle score (scaled), early-life lifestyle score (scaled), age (scaled), sex, and self-identified ancestry group (Senoi, Negrito, Proto Malay, multiple/other) as fixed effects, with a kinship matrix incorporated as a random effect. P-values were adjusted using the Benjamini–Hochberg false discovery rate (FDR) procedure [[47]](https://paperpile.com/c/zOgmfo/xwSh). Genes with FDR < 0.1 were considered significantly associated.

Pathway enrichment: Gene Ontology (GO) enrichment analyses were performed using clusterProfiler (v4.10.1) with GO Biological Process annotations from org.Hs.eg.db [[49,50]](https://paperpile.com/c/zOgmfo/wG7s+ob8C). Analyses were conducted separately for genes positively and negatively associated with current lifestyle. Because early life lifestyle had fewer total associated genes, and because biologically we expected early life environments to have bi-directional effects on immune processes, we grouped up- and down-regulated genes associated with the early life lifestyle index. For all analyses, the background set included all expressed genes tested in differential expression models. Enrichment was performed using enrichGO() with gene set size thresholds of 5–500 genes, and p-values were adjusted using the Benjamini–Hochberg procedure. GO term similarity and clustering were evaluated using pairwise_termsim(), and enriched categories were visualized with enrichplot.

Transcription factor enrichment: Transcription factor (TF) enrichment analyses were performed using TF–target interactions from the TRRUST v2 database [[51]](https://paperpile.com/c/zOgmfo/1Z7H). TFs with fewer than five target genes after filtering were excluded. Genes associated with lifestyle variables were grouped according to direction of effect (urban-associated, subsistence-associated, or early-life associated). To reduce redundancy among TFs with overlapping target sets, pairwise Jaccard similarity indices were calculated and hierarchical clustering was performed on the resulting distance matrix. TFs with highly overlapping target sets (Jaccard similarity ≥ 0.7) were collapsed into meta-TF clusters. Enrichment of TF target genes within each lifestyle-associated gene set was assessed using two-tailed Fisher’s exact tests against all expressed genes as background. P-values were adjusted using the Benjamini–Hochberg FDR procedure [[47]](https://paperpile.com/c/zOgmfo/xwSh), and TF clusters with FDR < 0.1 were considered significantly enriched.

TWAS enrichment analyses: We assessed whether genes associated with current or early-life lifestyle were enriched for genes implicated in complex disease traits from previously published probabilistic transcriptome-wide association study (PTWAS) datasets [[52]](https://paperpile.com/c/zOgmfo/sLsy). Gene-trait associations were obtained from the PTWAS multi-tissue resource, and Ensembl gene identifiers were converted to HGNC gene symbols using org.Hs.eg.db [[50]](https://paperpile.com/c/zOgmfo/ob8C). Analyses were restricted to noncommunicable disease traits that were selected a priori because these diseases differ substantially in prevalence between subsistence-level and market-integrated environments, resulting in 76 traits and 14,119 genes. Genes from our differential expression analyses were classified into three categories based on statistical significance (FDR < 0.1) and direction of effect: (i) genes positively associated with current lifestyle (“current market-integrated”), (ii) genes negatively associated with current lifestyle (“current subsistence-level”), and (iii) genes associated with early-life lifestyle regardless of effect direction. For each trait, enrichment of PTWAS-associated genes within each gene set was evaluated using Fisher’s exact tests. The background universe consisted of all expressed genes tested in the differential expression analyses and present in the PTWAS resource. Separate enrichment tests were performed for urban-associated, subsistence-associated, and early-life-associated gene sets. For each comparison, we calculated odds ratios, overlap counts, and p-values. Resulting p-values were corrected for multiple hypothesis testing using the Benjamini–Hochberg false discovery rate procedure.

KEGG pathway enrichment analysis: To evaluate whether genes associated with lifestyle exposures were enriched for T helper cell differentiation pathways, genes from the KEGG Th1 and Th2 cell differentiation pathway (KEGG pathway hsa04658) were retrieved using the KEGG REST API in R [[53]](https://paperpile.com/c/zOgmfo/ArSc). Gene symbols were extracted from the pathway annotation and matched to genes tested in the differential expression analyses. For each gene set of interest, enrichment of Th1/Th2 pathway genes was assessed using Fisher's exact test.

Assessment of individual lifestyle components: For analyses of how individual lifestyle facets are associated with transcriptional variation, separate mixed models were fit for each predictor variable, namely eosinophil percentage, physical activity (ENMO), market diet score, traditional diet score, body roundness index (BRI), log(IgE), and current lifestyle score. Only samples with data for all seven predictor variables were included in these analyses (n=660 samples). All predictors were z-score standardized prior to modeling for comparable effect sizes. All models controlled for early life lifestyle, self-identified ancestry group, age, sex, and genetic relatedness. To compare transcriptional signatures across predictors, pairwise correlations of gene-level effect sizes were calculated and visualized using hierarchical clustering and UpSet plots.

**Elastic net classifier**

Associated script: https://github.com/laylabrassington/ELvsCL/blob/main/scripts/modeling.Rmd

To assess whether transcriptomic profiles could predict lifestyle variation, we trained a regularized logistic regression model using elastic net penalization. Individuals were first categorized into lifestyle groups based on tertiles of early-life and current lifestyle scores, yielding four groups: individuals consistently living in subsistence-based, non-industrial environments (n=185), consistently living in urban, market-integrated environments (n=204), and individuals who transitioned between environments (non-industrial to urban, market-integrated (n=32) or urban, market-integrated to non-industrial (n=36)). Model training was restricted to individuals from the two consistent environmental exposure groups. Prior to model training, genes significantly associated with age or ancestry group were removed to reduce potential confounding effects (FDR < 0.1, n-genes = 5,158). Gene expression values for the remaining genes were used as predictors in a binomial elastic net regression model implemented using the glmnet package in R [[54]](https://paperpile.com/c/zOgmfo/AYco). Models were fit using a binomial error distribution with an elastic net mixing parameter of α = 1 (equivalent to LASSO regularization). The regularization parameter (λ) was selected by 50-fold cross-validation using the minimum cross-validation error criterion (lambda.min). Model performance was evaluated using leave-one-out cross-validation, in which each sample was iteratively withheld, the model refit on the remaining samples, and the held-out sample assigned a predicted probability of market-integrated classification. Predictive performance was assessed using receiver operating characteristic (ROC) analysis and area under the curve (AUC). A final model was then trained on consistent environmental exposure groups and applied to individuals from the remaining two transition categories.

Among individuals in the urban, market-integrated to non-industrial group, associations between time spent in the current community and elastic net model predictions were evaluated using self-reported duration of residence categorized as <1 year, 1–10 years, or >10 years. To assess whether longer residence in a subsistence-based, non-industrial environment was associated with model predictions (using a continuous assignment probability from 0 to 1), the relationship between ordered residence duration categories (scored as 1, 2, or 3) and model predictions was tested using Spearman’s rank correlation in the stats package in R [[55]](https://paperpile.com/c/zOgmfo/UUq8).

Supplementary Figures

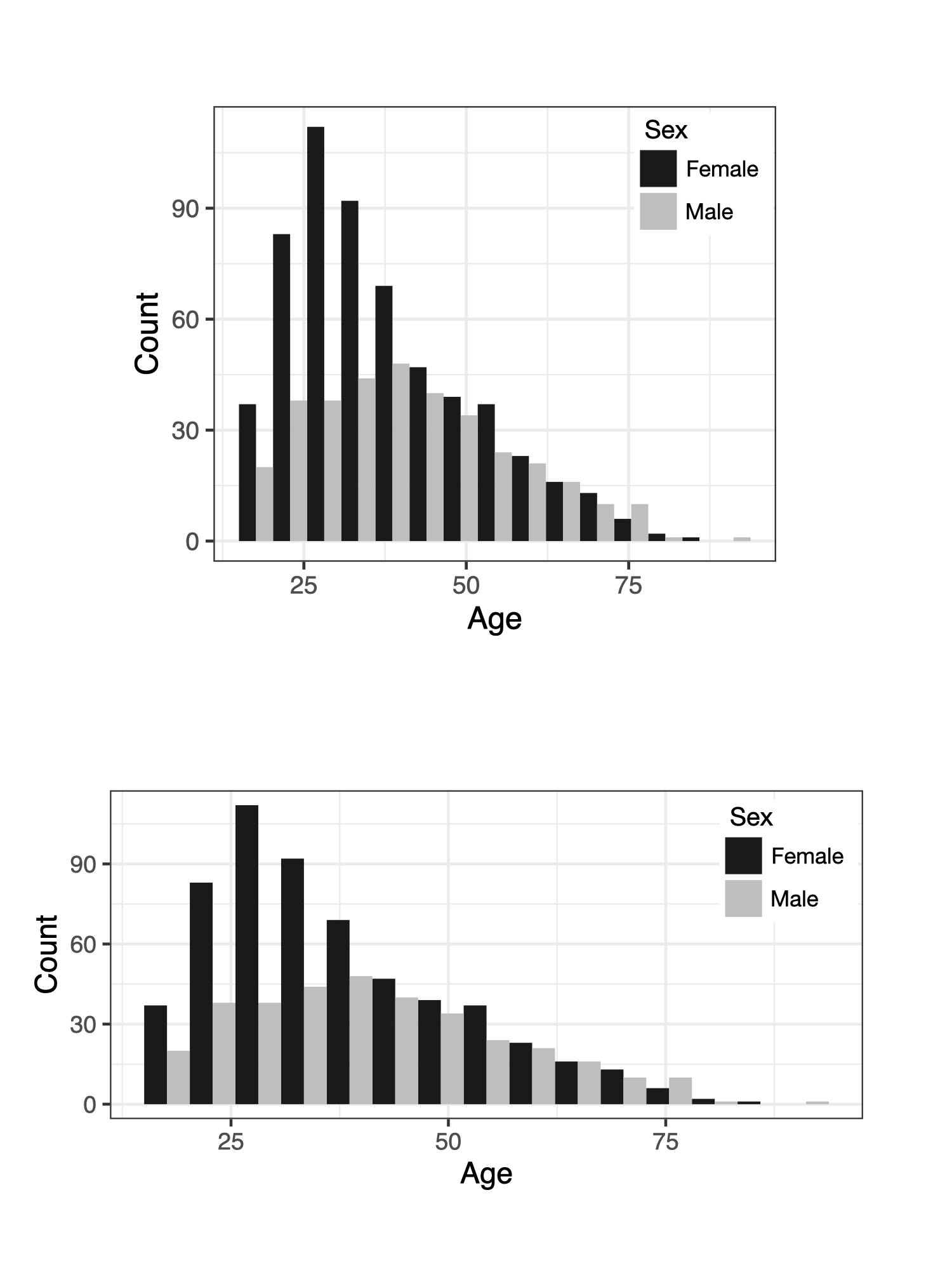

**Figure S1: Age and sex distribution of participants.** Participants included individuals from ages 18 to 91, and included 577 females and 345 males.

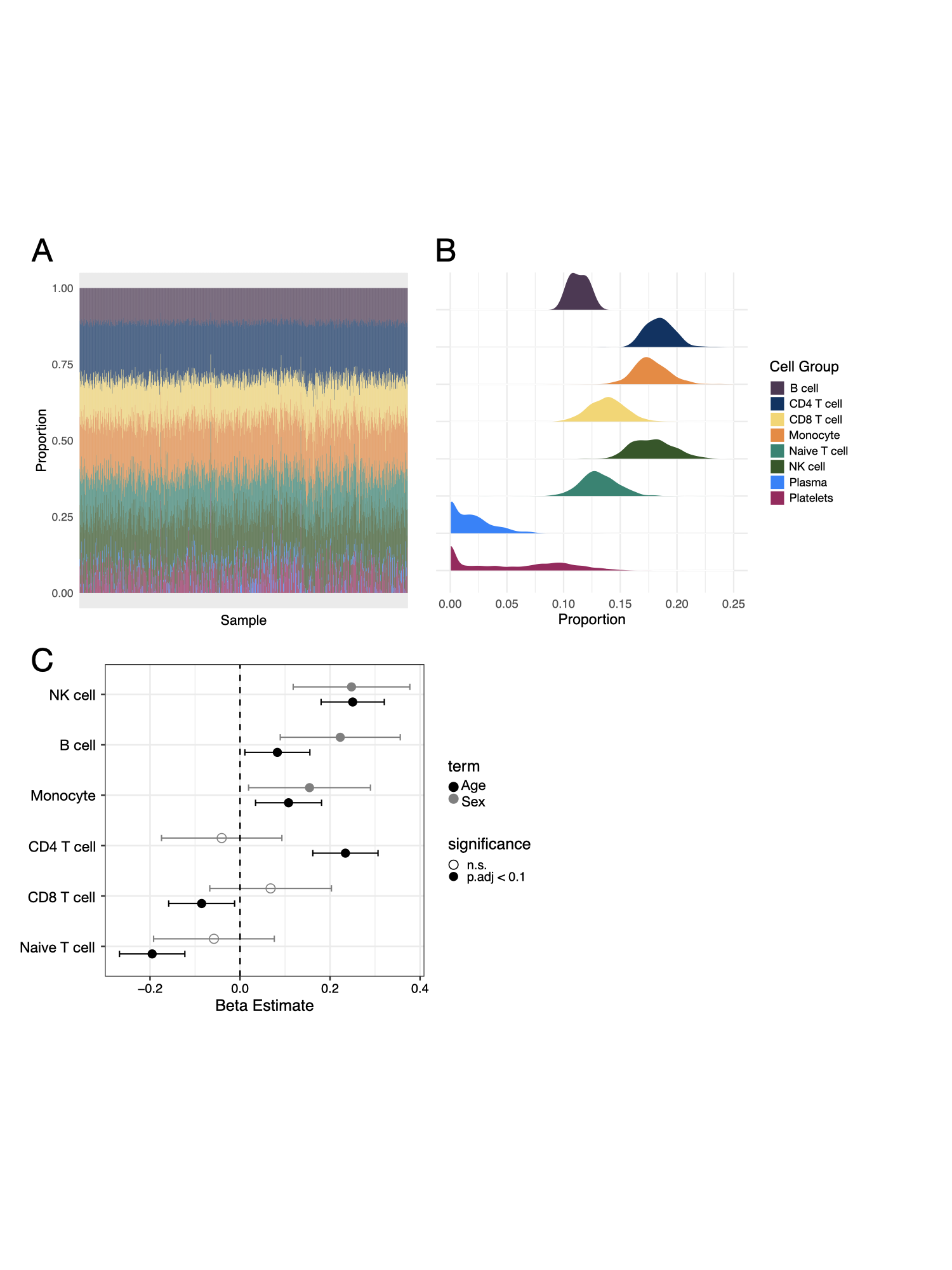

**Figure S2: Estimation of blood cell-type composition in Orang Asli individuals using an Orang Asli-specific single-cell RNA-seq reference panel.**

(A) Distribution of predicted blood cell-type proportions across all individuals.

(B) Distribution of predicted blood cell-type proportions within each individual, illustrating inter-individual variation in inferred cellular composition. Abbreviations: NK, natural killer.

(C) Associations between inferred cell-type proportions and age or sex estimated using linear models. Points indicate model effect estimates and horizontal bars indicate 95% confidence intervals. Filled symbols denote associations significant with an adjusted p < 0.1, whereas open symbols indicate non-significant associations.

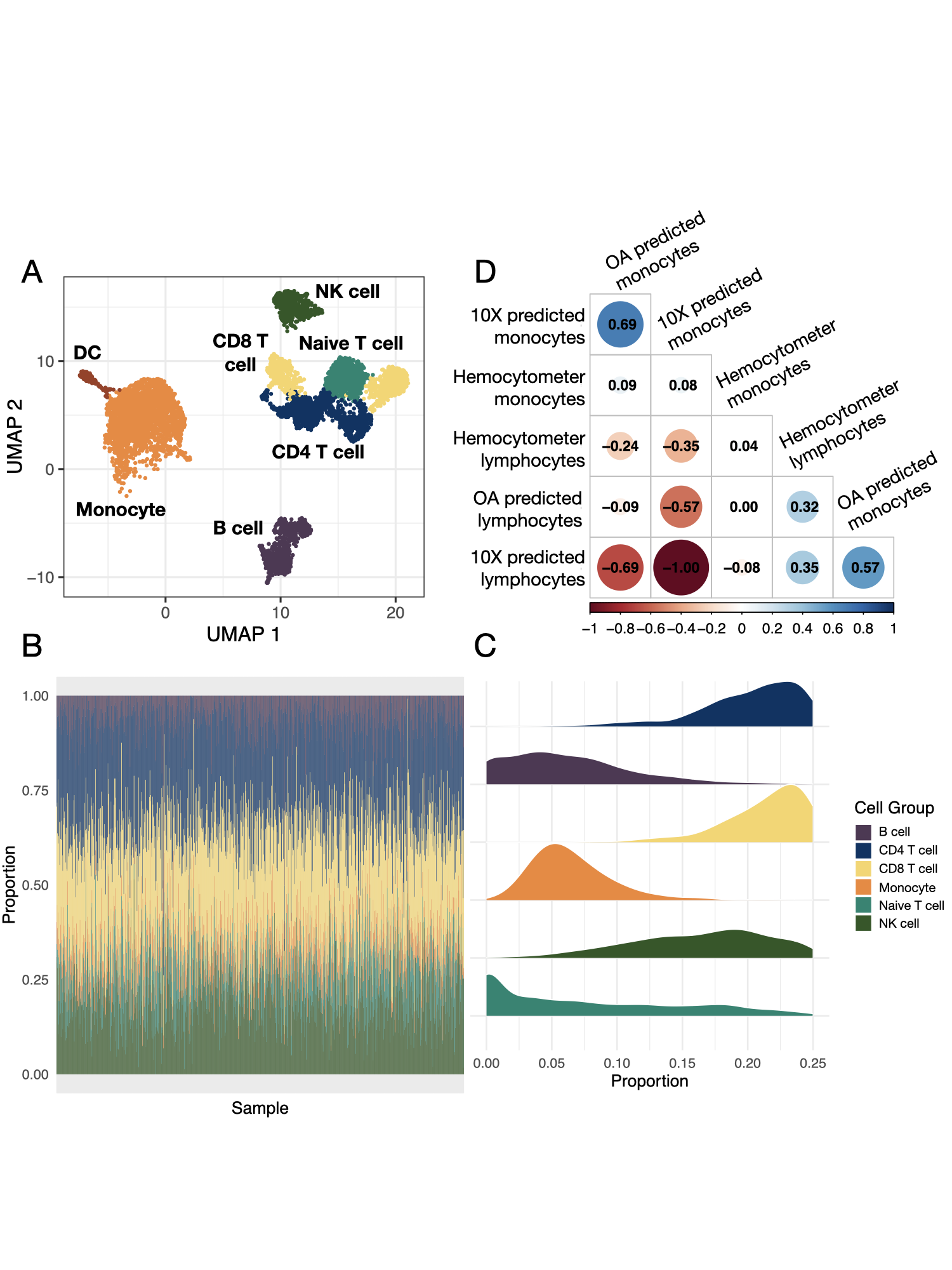

**Figure S3: Comparison of cell-type deconvolution results generated using a non-Orang Asli single-cell reference panel from 10x Genomics.**

(A) UMAP visualization of the non-Orang Asli single-cell RNA-seq reference dataset used for cell-type deconvolution. Abbreviations: DC, detritic cell; NK, natural killer.

(B) Distribution of predicted blood cell-type proportions across individuals inferred using the non-Orang Asli reference panel.

(C) Distribution of predicted blood cell-type proportions within individuals inferred using the non-Orang Asli reference panel.

(D) Correlation between cell-type proportions estimated using each reference panel and corresponding hemocytometer-derived cell counts.

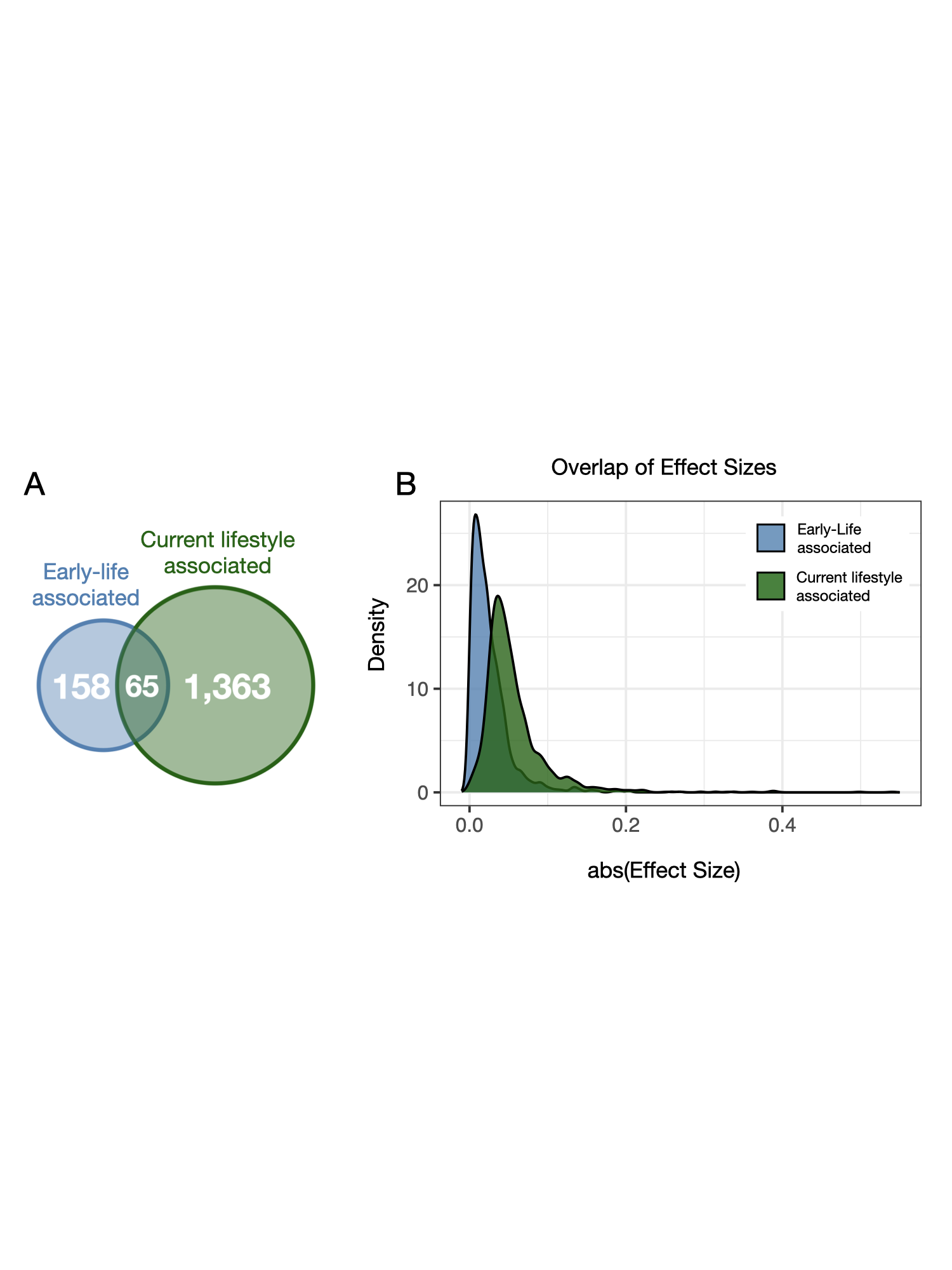

**Figure S4: Sharing and magnitude of early-life and current lifestyle effects on adult gene expression.**

(A) Venn diagram showing the number of genes significantly associated with early-life environment, current lifestyle, or both (adjusted p < 0.1).
(B) Distribution of absolute effect sizes for genes significantly associated with either early-life or current lifestyle environments (n = 1,586). Density curves summarize the magnitude of estimated effects, illustrating differences in the overall strength of transcriptional associations between environmental exposures.

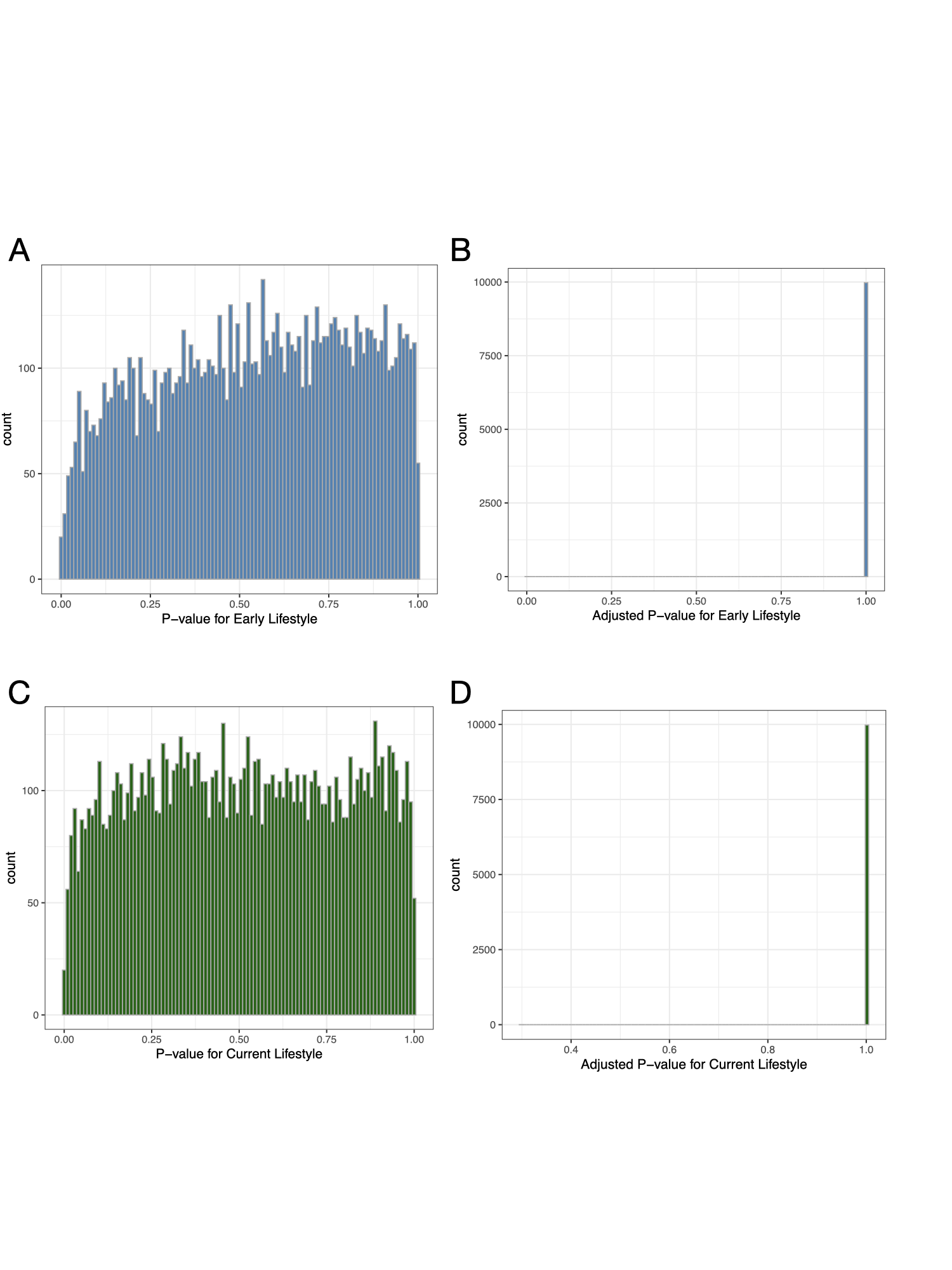

**Figure S5: Permutation analysis of early-life (EL) and current lifestyle (CL) gene expression model results.**

(A) Distribution of nominal p-values obtained for the EL model term across permutation analyses.

(B) Distribution of FDR-adjusted p-values obtained for the EL model term.

(C) Distribution of nominal p-values obtained for the CL model term.

(D) Distribution of FDR-adjusted p-values obtained for the CL model term.

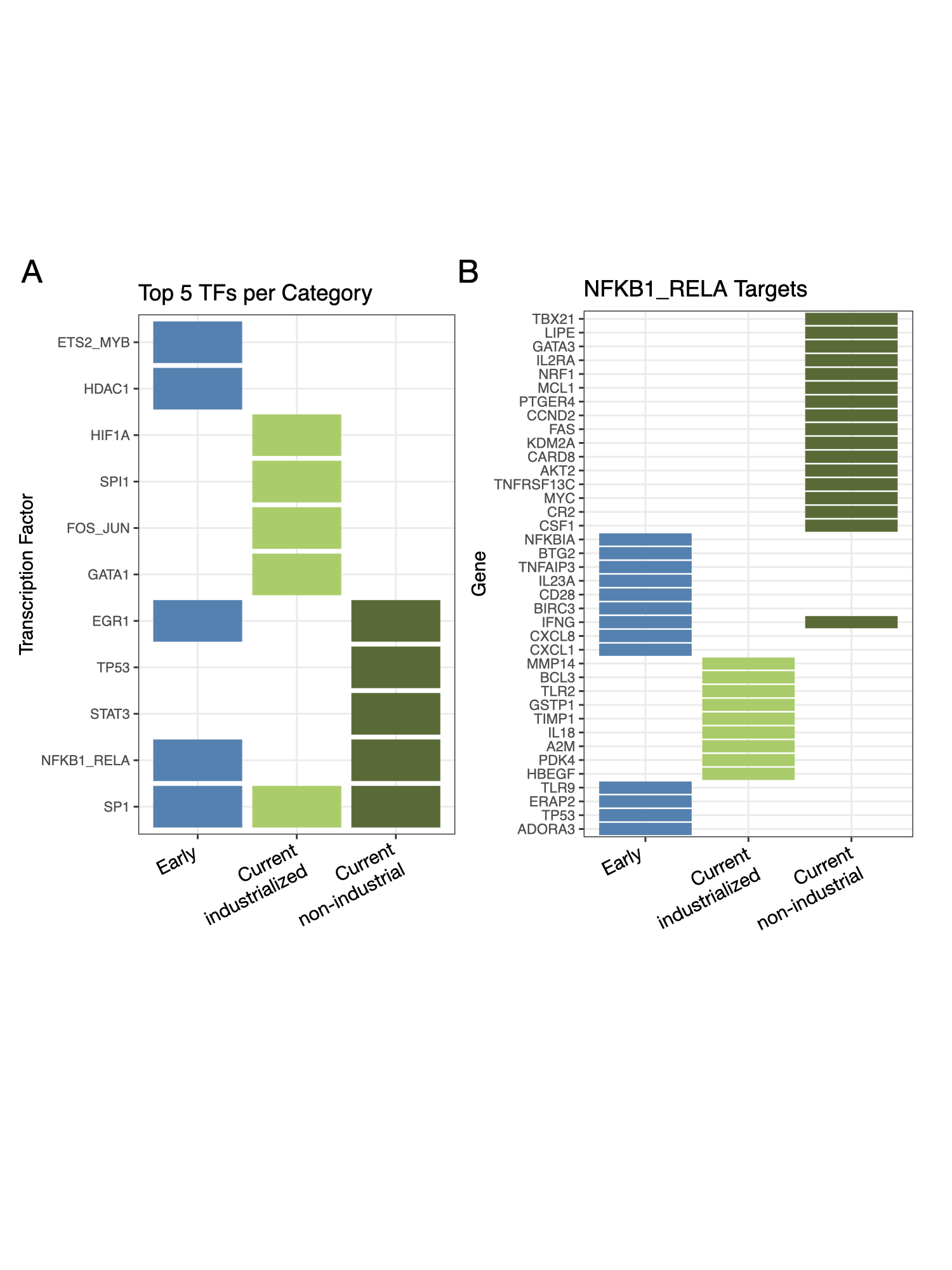

**Figure S6: Transcription factor target enrichment among lifestyle-associated gene sets.**

(A) Top five most significant transcription factor target enrichments for each gene category. Significance was assessed based on a Fisher's exact test of the overlap between lifestyle-associated genes and curated transcription factor target gene sets (see methods).
(B) Detailed overlap between targets of the NFKB1/RELA transcription factor complex and genes belonging to each lifestyle-associated gene category, highlighting shared genes contributing to enrichment signals.

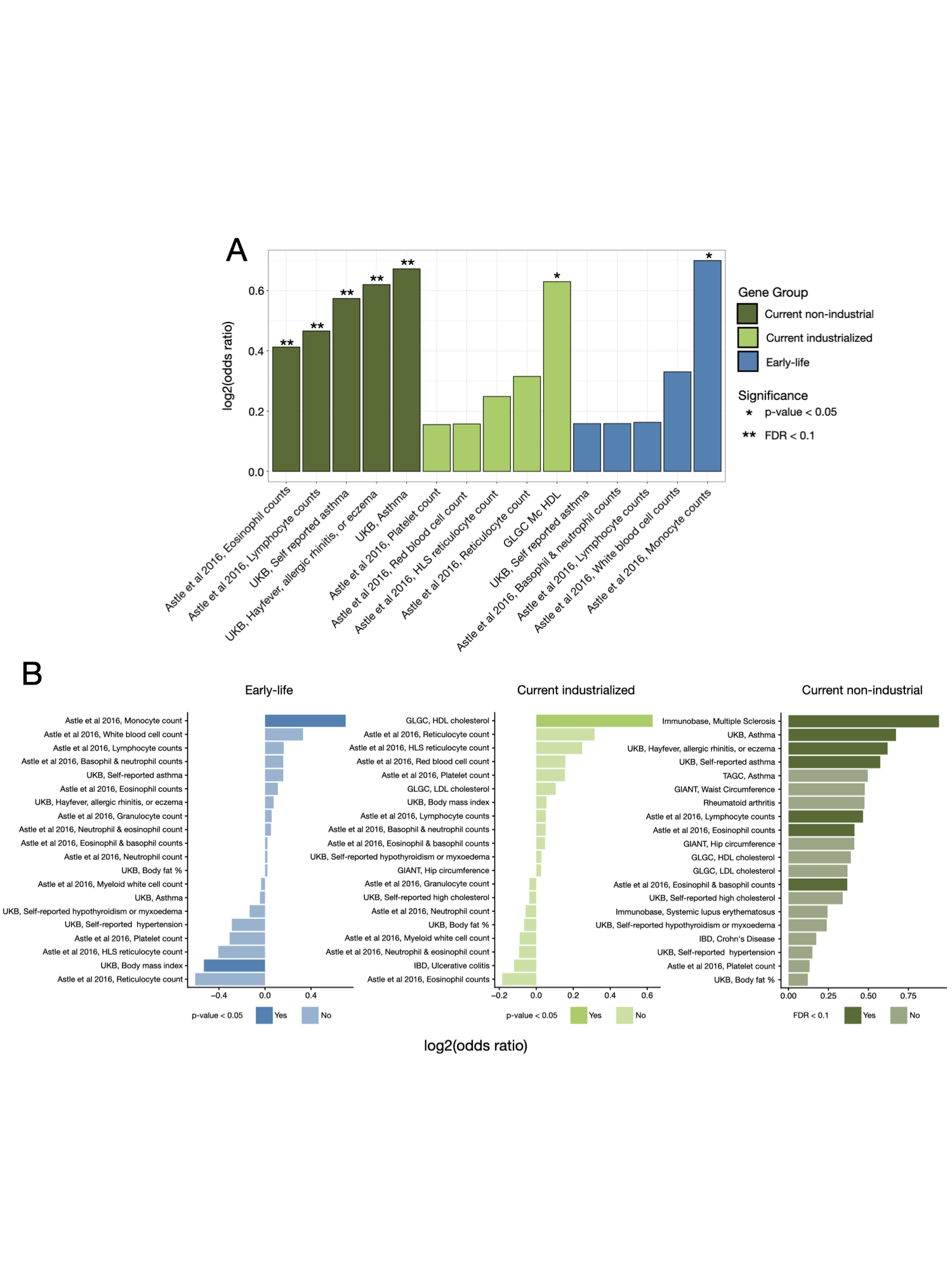

**Figure S7. Overlap between lifestyle-associated gene sets and transcriptome-wide association study (TWAS) results reported by Zhang et al. (2020).**

(A) Five most positively enriched traits for each gene group. Colors indicate gene group membership. Asterisks denote statistical significance (*p-value < 0.05; **FDR adjusted p < 0.1).
(B) Top 20 trait–gene group associations ranked by odds ratio. For the early-life and current industrialized gene groups, bar transparency reflects nominal significance, with more opaque bars indicating p-values < 0.05. For the current non-industrial gene group, bar transparency reflects FDR adjusted p-value significance, with more opaque bars indicating an FDR adjusted p-value < 0.1.

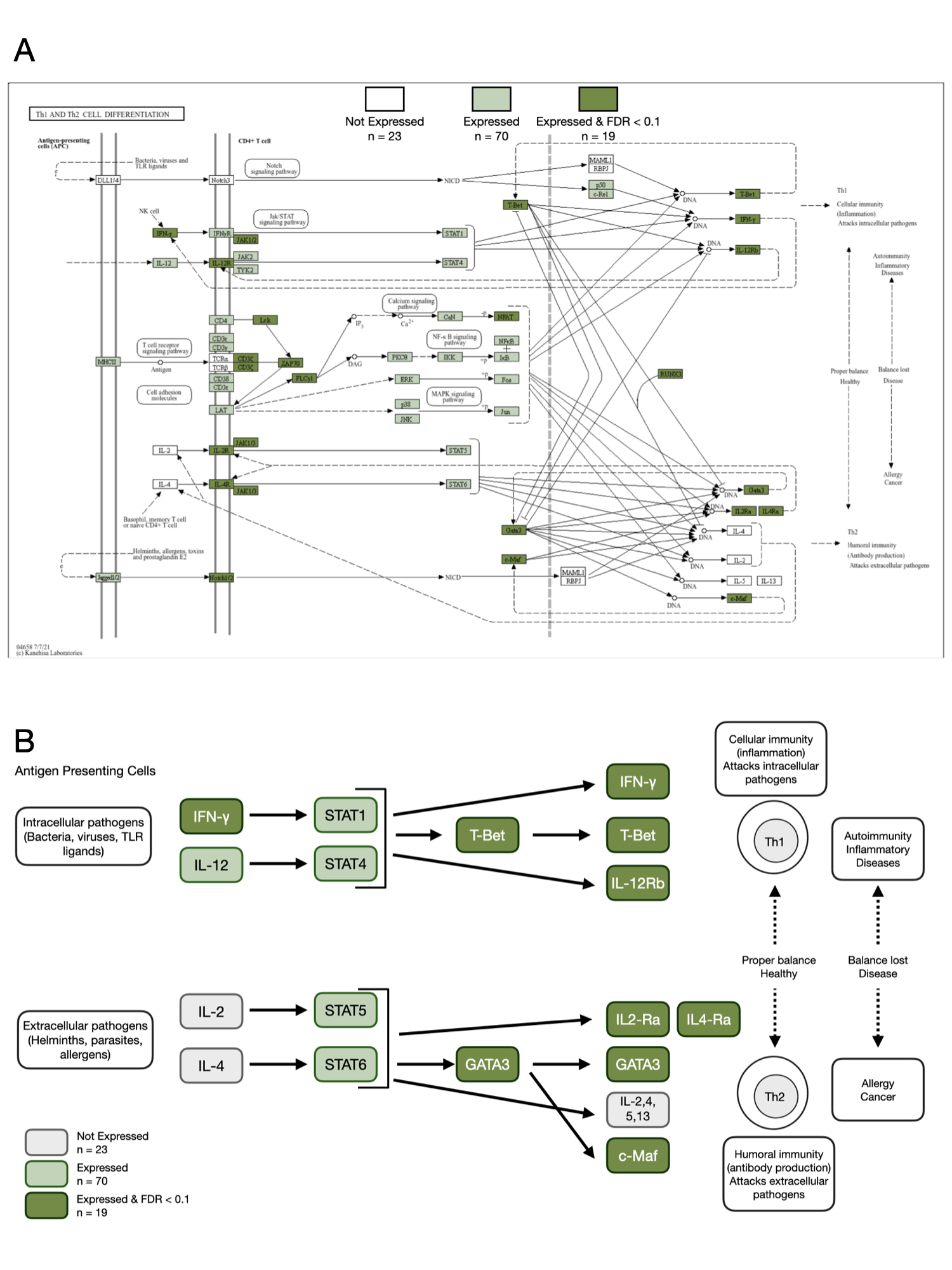

**Figure S8: Overlap between current subsistence-level genes and the T helper cell differentiation pathway.**

Full KEGG pathway diagram for T helper cell differentiation (hsa04658). Genes shown in white were present in the KEGG pathway but not detected in the study dataset. Genes shown in light green were detected but not significantly associated with current subsistence-level environments, whereas genes shown in dark green were detected and significantly associated with current non-industrial environments (FDR adjusted p < 0.1).

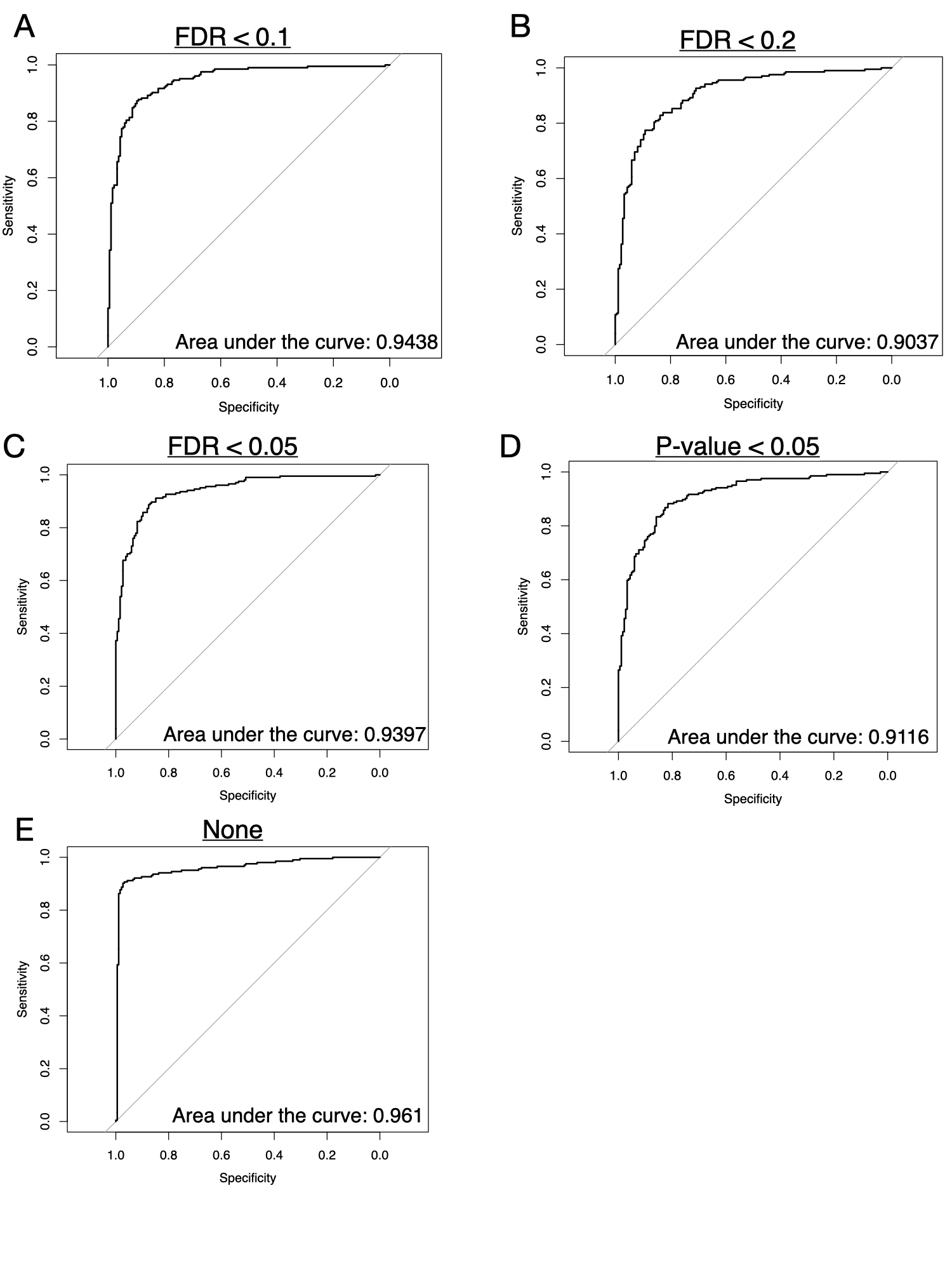

**Figure S9: Predictive performance of lifestyle classification models under alternative gene filtering criteria.** Area under the receiver operating characteristic curve (AUC) from leave-one-out cross-validation (LOOCV) for models trained using different gene inclusion/exclusion criteria based on FDR adjusted p-values. Comparisons assess the robustness of predictive performance to alternative approaches for controlling potential confounding effects of age and ancestry.

(A) Genes associated with age or ancestry removed at FDR < 0.1 (reported in main-text).

(B) Genes associated with age or ancestry removed at FDR < 0.2.

(C) Genes associated with age or ancestry removed at FDR < 0.05.

(D) Genes associated with age or ancestry removed at nominal P < 0.05.

(E) Model trained using the full gene set without filtering.

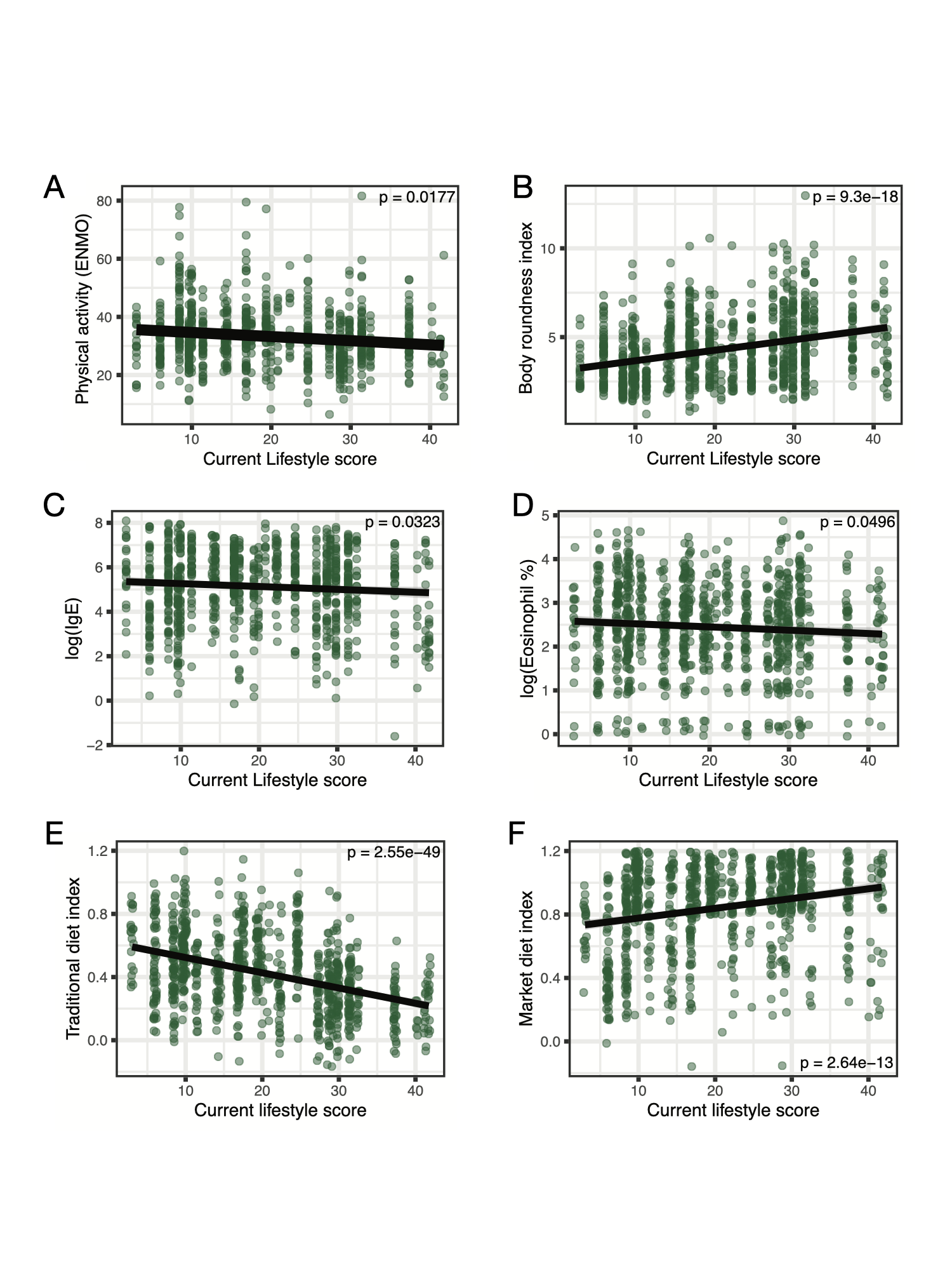

**Figure S10: Associations between current lifestyle score and four aspects of lifestyle relevant to immune function.**

(A) Association between current lifestyle and physical activity (ENMO).

(B) Association between current lifestyle and body roundness index.

(C) Association between current lifestyle and log(IgE values).

(D) Association between current lifestyle and log(Eosinophil %).

(E) Association between current lifestyle and a traditional diet index, as created by Watowich et al [[1]](https://paperpile.com/c/zOgmfo/vC7r), which in short, is the extent (or lack thereof) of traditional foods consumed.

(F) Association between current lifestyle and market diet index, as created by Watowich et al [[1]](https://paperpile.com/c/zOgmfo/vC7r), defined as the extent of market-derived foods incorporated in diet.
